# Non-Parametric Analysis of Inter-Individual Relations Using an Attention-Based Neural Network

**DOI:** 10.1101/2020.03.25.994764

**Authors:** Takashi Morita, Aru Toyoda, Seitaro Aisu, Akihisa Kaneko, Naoko Suda-Hashimoto, Ikuma Adachi, Ikki Matsuda, Hiroki Koda

## Abstract

1. Social network analysis, which has been widely adopted in animal studies over the past decade, enables the revelation of global characteristic patterns of animal social systems from pairwise inter-individual relations. Animal social networks are typically drawn based on geometric proximity and/or frequency of social behaviors (e.g., grooming), but the appropriate metric for inter-individual relationship is not clear, especially when prior knowledge on the species/data is limited.
2. In this study, researchers explored a non-parametric analysis of inter-individual relations using a neural network with the attention mechanism, which plays a central role in natural language processing. The high interpretability of the attention mechanism and flexibility of the entire neural network allow for automatic detection of inter-individual relations included in the raw data, without requiring prior knowledge/assumptions about what modes/types of relations are included in the data. For these case studies, three-dimensional location data collected from simulated agents and real Japanese macaques were analyzed.
3. The proposed method successfully recovered the latent relations behind the simulated data and discovered female-oriented relations in the real data, which are in accordance with previous generalizations about the macaque social structure.
4. The proposed method does not exploit any behavioral patterns that are particular to Japanese macaques, and researchers can use it for location data of other animals. The exibility of the neural network would also allow for its application to a wide variety of data with interacting components, such as vocal communication.

## 1 Introduction

Understanding the characteristics of animal social groups is a major objective of animal ecology. Complex social systems of eusocial insects and non-human primates are often viewed as biological models or precursors of human societies; as such, they have attracted the interest of social scientists as well. Social network analysis, which has been widely adopted in animal studies over the past decade, can be used to determine global characteristic patterns of animal social systems from pairwise inter-individual relations (Krause et al., 2015).

The application of social network analysis to animal studies requires an appropriate metric for the pairwise relationship between individuals (Sosa et al., 2020). Popular options for the relation metric include proximity, which is defined defined simply as geometric closeness between individuals (often thresholded and converted to counts of co-appearance within the threshold; Silk et al., 2006b,a; Croft et al., 2008; Clark, 2011; Boyland et al., 2013; Castles et al., 2014; Krause et al., 2015; Schofield et al., 2019; Gelardi et al., 2020), and frequency of social behaviors (e.g., grooming of non-human primates; Chepko-Sade et al., 1989; Silk et al., 2006b,a; Croft et al., 2008; Clark, 2011; King et al., 2011b,a; Castles et al., 2014; Krause et al., 2015). However, an appropriate relation metric is not necessarily clear, especially when prior knowledge about the species/data is limited (cf. Haddadi et al., 2011; Castles et al., 2014; Farine and Whitehead, 2015; Canteloup et al., 2020). For example, location data for measuring the geometric proximity is available for a wide variety of animals. This includes species whose social activities are not yet well-known. Location information has become more reliable and scalable thanks to recent developments in bio-logging technology, such as GPS (Heupel et al., 2006; Rutz and Hays, 2009; Rutz et al., 2012; Fehlmann and King, 2016; King et al., 2018; Dore et al., 2020; Gelardi et al., 2020). However, naively measuring the Euclidean distance between individuals and treating pairs with a smaller distance as socially related is not necessarily reasonable. Some animal agents, for instance, may follow other individuals while keeping a certain geometric distance (perhaps to avoid being attacked), but such distant relationship/dependency would be missed if only proximal co-occurrences are counted as the index. Distance could also have different social meanings if other factors, such as trees and other objects, intervene (Pays et al., 2007). Furthermore, vertical divergences are expected to be less or irrelevant in an analysis of terrestrial animals, which do not fly or dive, so cannot control their vertical distance from other individuals.

Given the limitations of the conventional analytical paradigm depending on manually selected relation metrics, the present study explores a non-parametric, data-driven analyzer of inter-individual relations. Unlike conventional analysis, the proposed approach automatically detects inter-individual relations from raw data using a machine learning technique (Figure 1; cf. Maekawa et al., 2020). For example, location logs of animals collected through bio-logging devices can be analyzed without the need to first determine relevant aspects of the data such as co-appearance within a certain distance. The proposed method acts as a starting point for a series of data analyses (highlighted in yellow in Figure 1), and the detected relations can be used in additional analyses, including modes/types of relations and social networks. Under the proposed framework, geometric proximity would be reassessed as an instance of such post-hoc analyses, rather than the definition of relationship.

**Figure 1:**
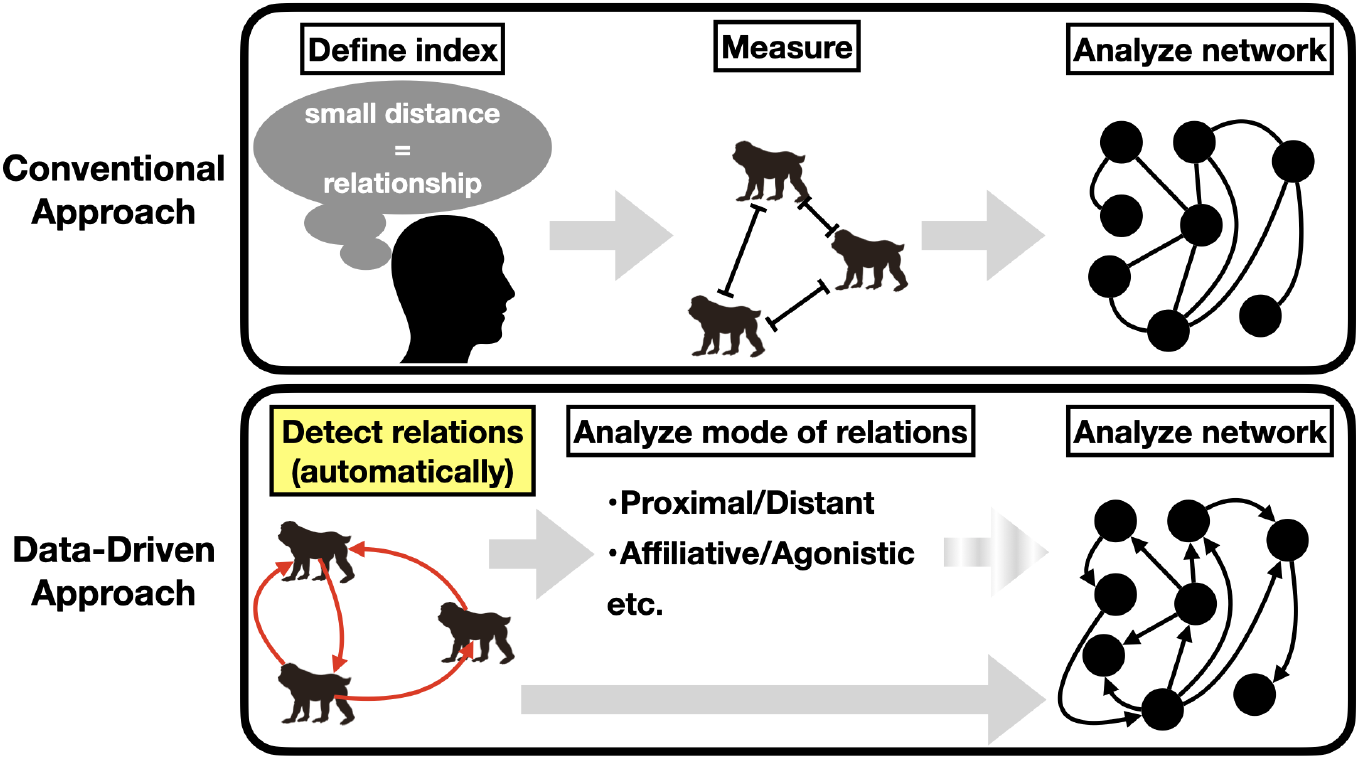
Comparison of the conventional approach to analysis of animal social relations/systems (top) with the alternative data-driven approach proposed in the current study (bottom). Using the conventional approach, researchers first need to define an appropriate index for measuring relationship between individual animal agents, which can be difficult for understudied animals. By contrast, the proposed approach first detects inter-individual relations automatically by using machine learning techniques, without manual specification of relevant aspects of the data, and then uses the detected relations to further characterize the modes/types of relations and more global patterns via social network analysis. The present study proposes a method for the first step of the data-driven approach (highlighted in yellow).

The proposed non-parametric analysis of inter-individual relations is based on an artificial neural network, which is the canonical framework in today’s machine learning. Neural networks are universal approximators of mathematical functions (i.e., they can approximate *any* types of computable functions, if ideally tuned; Cybenko, 1989; Hornik, 1991; Jin et al., 1995; Perez et al., 2019) and, thus, can be used for analysis of a broad range of data without assumptions about random distributions and latent structures behind the data. One drawback of using neural networks, however, is that they are often non-interpretable. Neural network computation consists of thousands of summations, multiplications, non-linear activations (e.g., sigmoid and tanh), and other basic calculations, and does not explain what aspects/portions of data are important. To perform interpretable analysis of inter-individual relations, the present study exploits a special neural network module called the *attention mechanism.* The attention mechanism was originally invented for machine translation to represent relations between words in the source and target languages (Figure 2a; Bahdanau et al., 2015). More recently, studies have used the attention mechanism to represent word relations within input sentences (self-attention, Figure 2b; Vaswani et al., 2017; Devlin et al., 2018). It is now considered central technology in today’s natural language processing (NLP). The present study applies this technology to reveal latent relations among animal agents that are reflected in their location data, without prior assumptions about the possible types of relationship structures or noise distributions (Figure 2c; see Yuan and Jia, 2019; Maekawa et al., 2020, for other applications to biological studies). While only the location data are discussed in this study, there are many possible applications of our relation analysis given the flexibility of the neural network.

**Figure 2:**
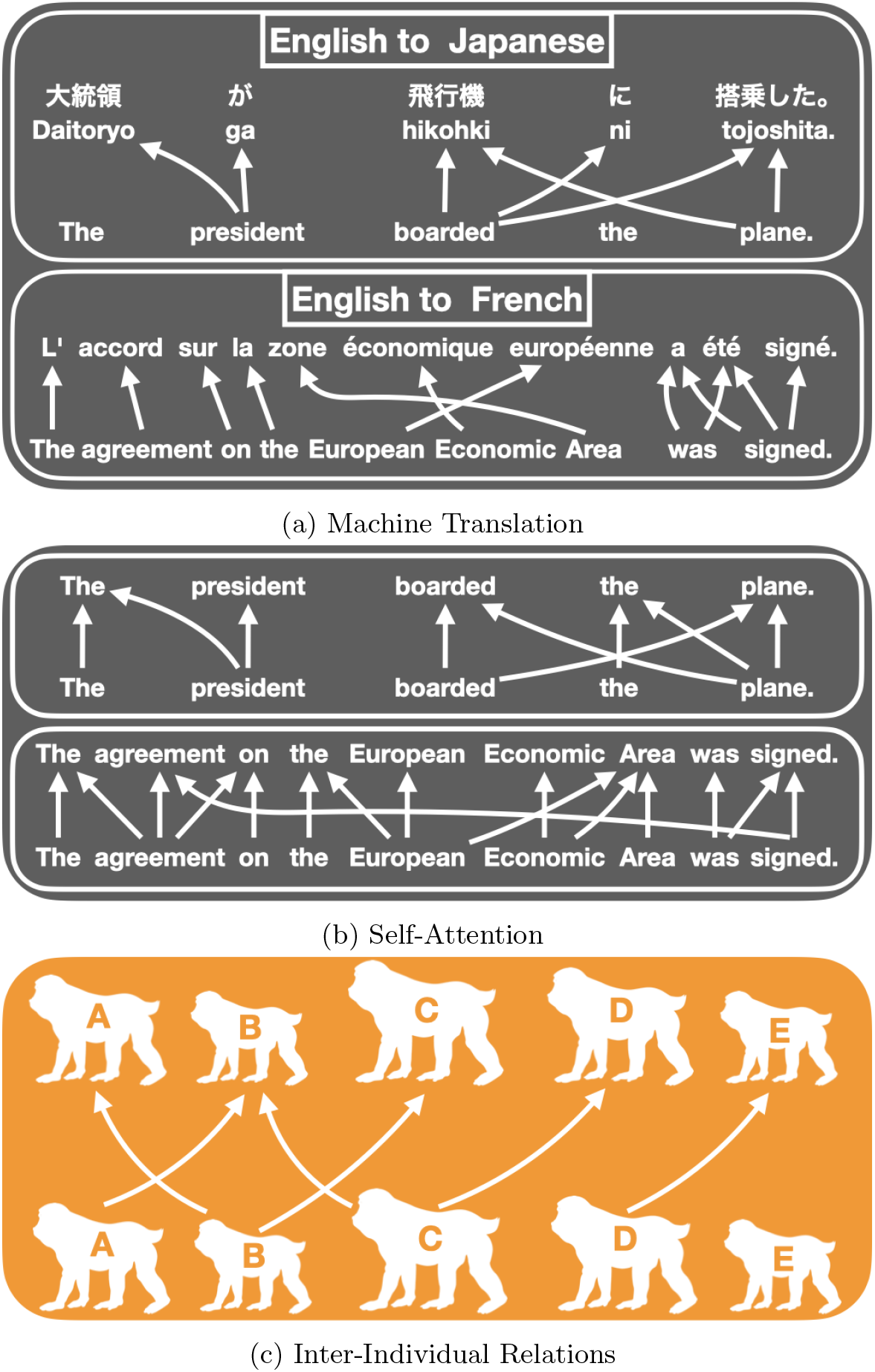
History of the use of the attention mechanism. (a) The original use of the attention mechanism in machine translation, relating words in the source language (English) to ones in the target language (Japanese/French). (b) More recent application in NLP that represents relations between words within the input sentences. (c) Application to inter-individual relations explored in this study.

The proposed method is explored using three-dimensional (3D) location data collected from Japanese macaques *(Macaca fuscata*) as well as simulation data that clarify its advantages over conventional distancebased analysis. Japanese macaques show stable relationships structured on dominance and kinship, and they have been studied extensively for more than 50 years to understand the evolution of social systems in primate lineages, including humans (Nakagawa et al., 2010). This large accumulation of knowledge and data makes the Japanese macaque an ideal species for evaluating our novel analysis. We emphasize that this method is not limited to this species or primates generally, and could even be more valuable for studying animals that do not show interpretable social activities.

## 2 Materials & Methods

### 2.1 Attention-Based Analysis of Inter-Individual Relations

This section provides an overview of the proposed methods in §2.1.1 followed by a more detailed explanation in §2.1.2–2.1.3.

#### 2.1.1 Overview of the Methods

The framework of the relation analysis in this study is to (i) train a neural network that predicts the 3D location of each individual macaque—simulated or real—from the locations of other individuals, and then (ii) check how much attention the network pays to each of the referent individuals when it makes the predictions. For simplicity, the network receives the location data at a single time point only and does not analyze temporal changes in the data, although the temporal information could include important traits of individuals and groups (cf. Morita et al., 2020) and thus should be discussed in future studies.

Figure 3 shows a sketch of the network structure, without details. The location of an individual named *A* is predicted from that of four other individuals, *B, C, D,* and *E.* The prediction is made from the weighted sum of the (transformed) location information of the four referents, where the relevance of each referent to *A—*called *attention weight—*is also derived from the location information. The attention weights govern the information flow from the referents; they intuitively represent *who is contributing how much to the prediction*. Our analysis of the relations between the target and referents is performed considering these attention weights.

**Figure 3:**
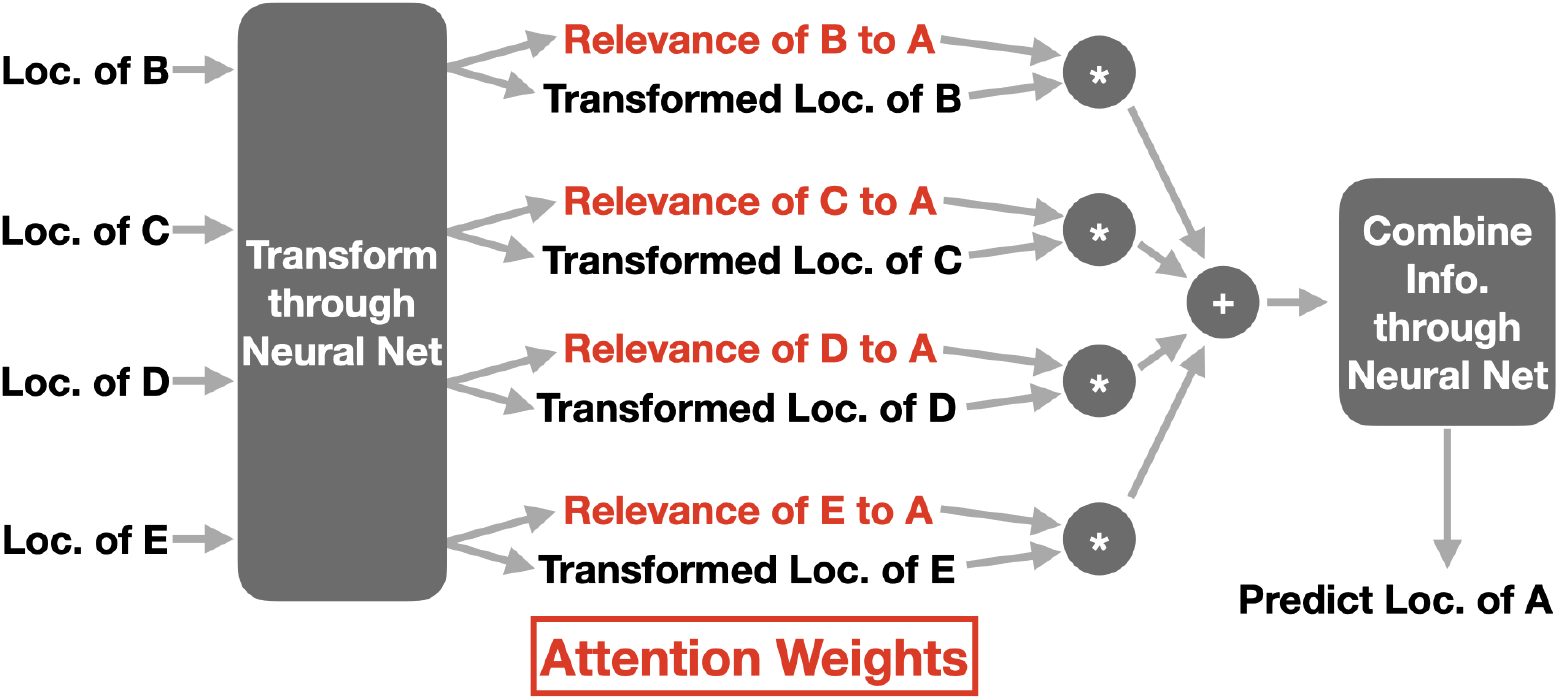
Schematic diagram of attention-based analysis.

The attention weights were normalized and always have a sum of 1. An inherent disadvantage of this formalization is that the model never detects the independence of the target individual, as at least one referent will have a non-zero weight. To get around this, the predictive performance of the attention model was compared with a baseline model that predicts the target without reference to other individuals. If the attention weights are redundant, the attention model will have the same level of predictive performance as the no-referent baseline. The baseline model was trained to maximize the likelihood.^1^

#### 2.1.2 Detailed Description of the Neural Network Structure

This section provides a detailed description of the attention-based neural network used in this study. As explained in the previous section, the neural network predicted the discretized location *y_i_* ∈ {1,…,*C*} (*C* = 400) of each individual *i* ∈ {1,…, *N*} from the continuous 3D location of the other *N* – 1 individuals, **x**_1_,…, **x**_*i*-1_, **x**_*i*+1_,…, **x**_*N*_ (**x**_*j*_ ∈ ℝ^3^). The predictions were made in the form of conditional probability: that is, 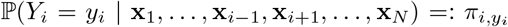. To efficiently compute all the combinations of the target and referent individuals, the researchers customized Transformer (Vaswani et al., 2017; Devlin et al., 2018) to fit the study’s purpose (i.e., the analysis of inter-individual relations), preventing the undesirable flow of information (e.g., the location of an individual should not be used when predicting that individual’s own location). This allowed us to represent the input (continuous 3D location) and output (conditional probability of each possible discretized location) of all individuals in each single matrix, which can be expressed as **X**:= (**x**_1_,…, **x**_*N*_)^*T*^ and **Π**:= (*π_i,c_*)_*i*∈{1,…,*N*},*c*∈{1,…,*C*}_ respectively. The **x**_*j*_ was scaled (by the upper bound in each dimension, see §2.2 for the bounds) and shifted so that each dimension ranged between – 1 and 1.

Figure 4 shows the network architecture, which is composed of two attention layers. Attention layers are a neural network module that take two types of inputs: *query* (**Q**) and *memory* (**M**). In the current study, **Q** and **M** were matrices of the same size whose rows represented the *N* individuals and whose columns represented the arbitrary number of dimensions in hidden layers. The memory encoded the location information of the referent individuals. The query, on the other hand, was either a vector-represented identity information of the target individuals (first layer) or the output from the first attention layer (second layer). The memory was linearly transformed into two matrices of the same size, called *key* (**K**) and *value* (**V**; see Figure 5b). The core computation in the attention layers was a weighted summation of the row vectors of **V**, which corresponds in this study to combining the information about the referent individuals proportional to their importance. The weights for this summation (i.e., the attention weights) were computed from the dot product of the row vectors of the key and query; equivalently, matrix multiplication **KQ**^T^ was computed. The attention weights were obtained by scaling the dot product by the square root of the hidden dimensionality (i.e., number of columns in the key and query) and normalizing the outcome so that the sum over the referent individuals (i.e., rows of **KQ**^T^) was 1.0.

**Figure 4:**
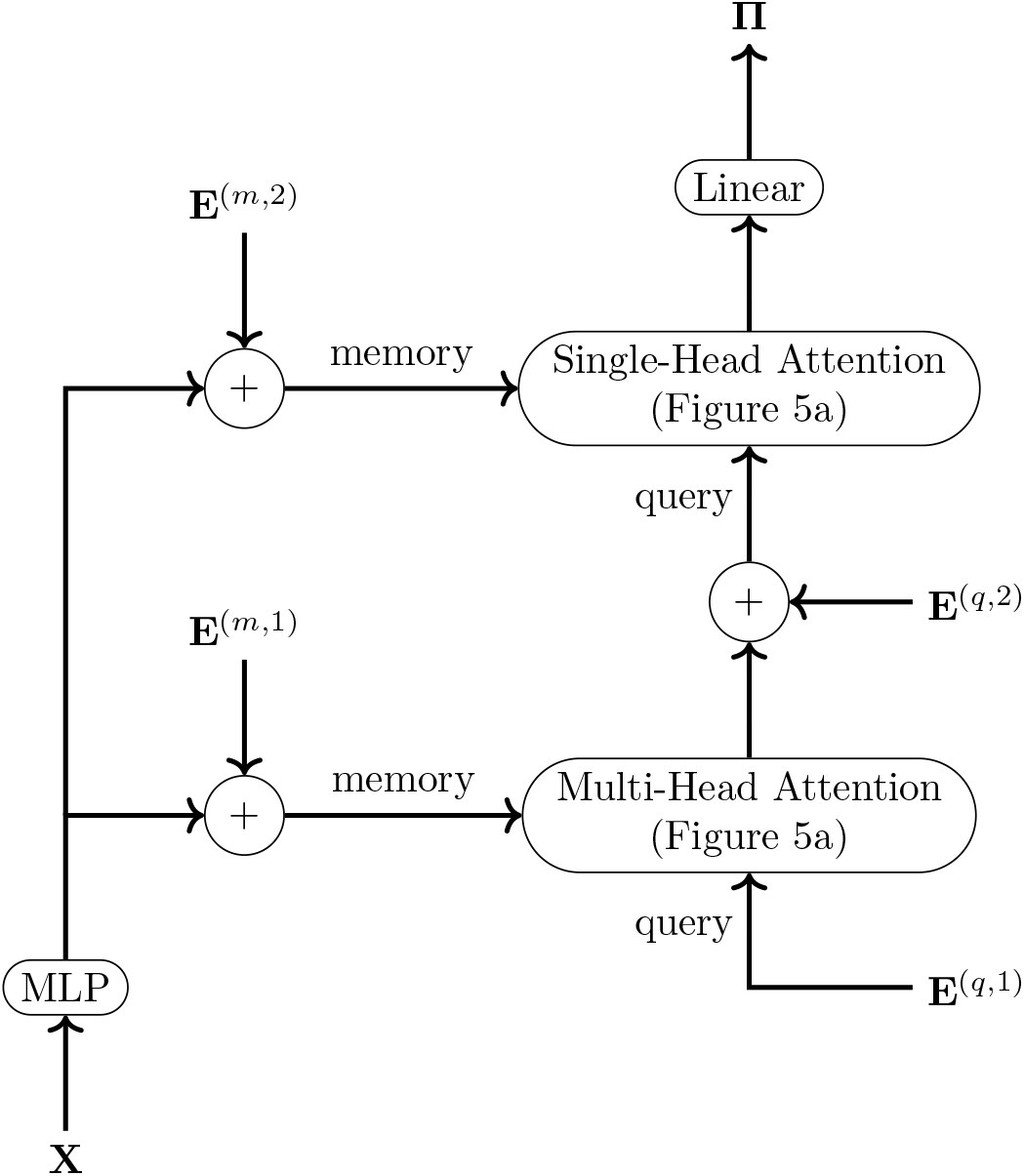
The entire structure of the neural network.

**Figure 5:**
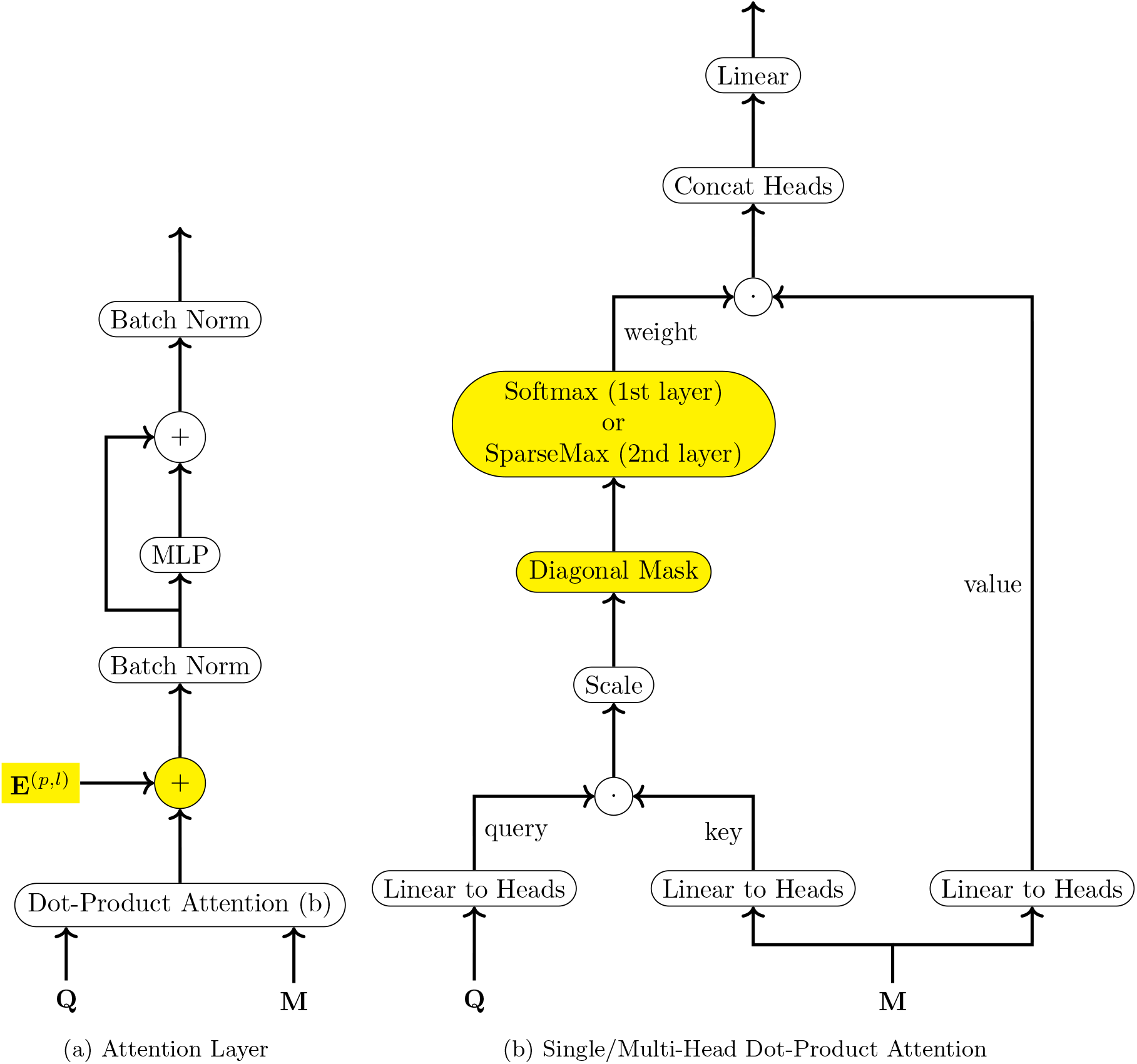
Structure of the attention layers (a) and their single/multi-head dot-product attention module (b). The nodes highlighted in yellow include modifications to the canonical design in Transformer.

In NLP applications, attention layers are usually split into several computations in parallel, called *heads.* Those multiple heads are implemented simply by splitting the hidden dimensions of the query, key, and value after the first linear transform (i.e., each head performs the computation using their submatrices grouped by columns; see Figure 5b).^2^ Having multiple heads often improves model performance because different heads can have different attention weights and, thus, different roles. In machine translation, for example, some heads may concentrate their attention weights on a single word in the source language upon the prediction of the corresponding word in the target language, while other heads may distribute their attention weights more broadly to incorporate the context of the source language sentences. In the current study, the first and second attention layers had different numbers of heads owing to their different roles. On one hand, the second layer was single-headed, and its attention weights were used to analyze inter-individual relations; this simpler architecture made interpretation of the attention weights easier. On the other hand, the role of the first layer was to provide useful information for the second layer (see the next paragraph for more details), and it was not used for an analysis of inter-individual relations. Thus, multiple heads were implemented in the first layer to improve the model performance of the training objective.

The major difference between this study and standard applications of Transformer/self-attention is that here, the network only uses the input location data as the *memory* of the attention. If we also use the input location for the *query,* then the network would be notified about which individual is geometrically closest to the target (as the attention weights are given by the similarity between the query and the key, which is computed from memory) and may simply use the closest individual for prediction. This is not the intended interaction of the investigation. Instead, in the first attention layer, the query only contained information about *whose location was being predicted as the target,* which was represented by trainable individual embeddings **E**^(*q*,1)^ (cf. akin to the time encoding in Shaw et al., 2018; Dai et al., 2019). The second layer received output from the first layer—in which the location of each referent individual (excluding the target) was encoded in a single vector per target—and used it as its query, together with other individual embeddings **E**^(*q*,2)^. The second layer thus computed the attention weight of each referent individual from three sets of information: (i) the individual’s location (memory), (ii) the location of the other referents (query, output from the first layer), and (iii) the name of the target individual being predicted (query), while the first layer had access only to (i) and (iii).

Note that the input location vector **x**_*i*_ and its transformed representation after the multilayer perceptron (MLP) did not encode *who* was located in that location (i.e., the network did not see the “*i*”, the row index in the matrix **X**; cf., the necessity of time-encoding in Transformer). This is why we added the individual embeddings **E**^(*m*,1)^ and **E**^(*m*,2)^ to the memory.

Other customizations inside the two attention layers are highlighted in yellow in Figure 5. First, we masked the diagonal entries of the attention weight matrix (Figure 5b); otherwise, the network allows for redundant predictions whereby a target individual’s location could be identified from its own input value. Secondly, the softmax function was replaced with the *sparsemax* (Martins and Astudillo, 2016; Niculae and Blondel, 2017; Niculae et al., 2018) to compute the attention weights in the second layer (Figure 5b), where the analysis of inter-individual relations was performed. This change made the analysis more robust and interpretable, because the sparsemax function can assign zeros to unmasked entries and express “no contribution”, whereas the softmax function always assigns non-zero weights. The softmax was kept in the first layer to create a single-vector representation of *all* the referent individuals, and the non-zero weights assigned by the softmax function were determined to be appropriate. Finally, the residual connection around the dot-product attention was removed (Figure 5a. The other residual connection around MLP was kept) because, aside from the information gated by the attention module, the residual connection also leaks information about the referent location, which makes the attention weights uninterpretable. Instead of a residual connection, individual embeddings **E**^(*p,l*)^ (where *l* ∈ {1, 2} denotes the layer) were added to the output from the dot-product attention. Since we masked the diagonal entries of the attention weight matrix, the output lost information about who was being predicted. Hence, we needed to retrieve that information before the final predictions could be made based on the output from the second attention layer.^3^

All hidden units in the network had 512 dimensions, except that the dimensions were evenly split among the four heads (128 dimensions each) in the attention layer.

#### 2.1.3 Training Procedure

The network was trained for 30, 000 iterations. Researchers used the Adam optimizer with a learning rate of 0.005, *β*_1_ = 0.9, *β*_2_ = 0.999, and a weight decay of 0.01. The learning rate was updated based on Transformer studies (Vaswani et al., 2017; Devlin et al., 2018), with 3,000 warm-up iterations. The batch size was 512. Dropout was applied at a rate of 0.1 to the output from the MLPs (Figure 4,5a) and the dot-product attention (Figure 5a before the addition of **E**^(*p,l*)^), but was not applied to the attention weights. This is because the number of attendees was small (five) in this study and robust inference is needed for that part.

Using one NVIDIA RTX 2080Ti graphic card, it took approximately nineteen hours to train the model on a five-agent data. It took longer to analyze more agents and the fifty-agent simulations took 148 hours to process.

### 2.2 Data

Attention-based relation analysis was performed on several simulated datasets (§2.2.1) and one real dataset (§2.2.2).

#### 2.2.1 Simulated Data

The reliability of the attention-based relation analysis was tested on several types of simulated data. Following the real data collected from Japanese macaques (see §2.2.2), both simulations had five individuals located in bounded 3D space ([0.0, 5.0] × [0.0, 4.0] × [0.0, 2.5]), which resembled the group cage where the macaques were kept.

The simulations consisted of two factors. The first factor governed relational structures of the animal agents. One of the structures modeled the scenario in which the five agents moved in a sequence, with each one following another (Figure 6a). Another structure modeled the hub and spoke relation in which a single hub individual was followed by the other four spoke dependents (Figure 6b). Besides these manually defined patterns, nineteen other relation structures were studied. Nine of the relation structures exhausted the patterns wherein each agent followed at most one precedence and no agent was isolated (i.e., the directed non-isolated acyclic graphs whose maximum indegree was one, modulo isomorphism). The other ten, whose maximal degree was two or greater, were randomly selected (see the supporting information S1.1 for details on the random sampling procedure).

**Figure 6:**
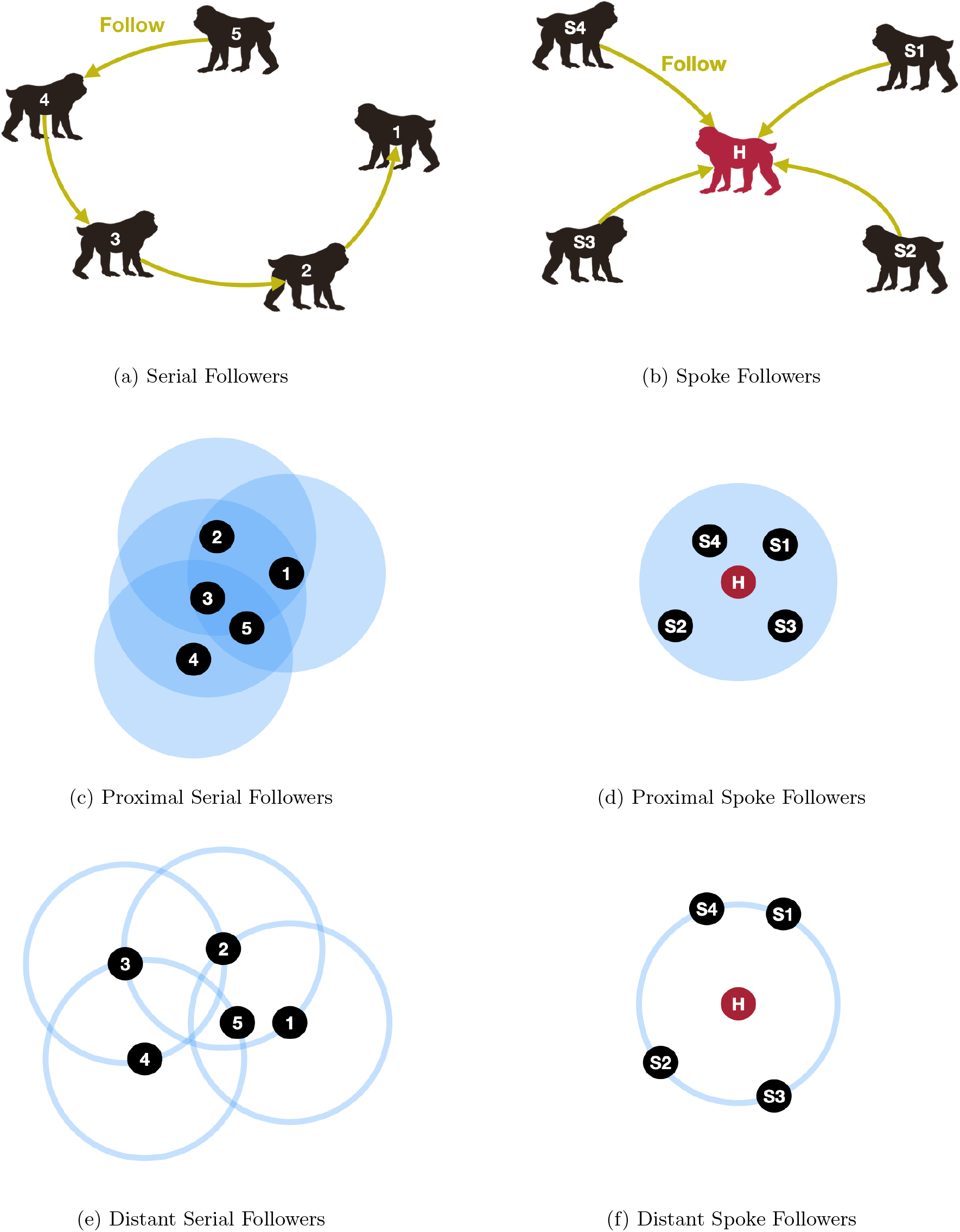
Two patterns of simulated data. (a) and (c) model animal agents that move in a sequence. (b) and (d) model the situation where there is a single hub individual (red) that is followed by the other four spoke dependents (black). (a) and (b) depict the relations among the animal agents modeled in the simulations. (c) and (d) describe the distributions from which the location data are sampled: The blue circles represent the support of the uniform distribution centered at each following individual.

The second factor parameterized pairwise dependency patterns between the related agents. One of the dependence patterns modeled a proximal situation in which each follower agent was always positioned around its precedent agent. Specifically, the serial-follower condition located the *(i* + 1)-th individual uniformly at random in the intersection of the domain and the sphere with a radius of 1.5 centered at the i-th individual’s location, which was implemented by rejection sampling (Figure 6c). Similarly, the hub-spoke condition sampled the location of the spoke individuals uniformly from the intersection of the domain and the sphere with a radius of 1.5 centered at the location of the hub individual’s location (Figure 6d). By contrast, the other dependence pattern modeled a less close relation in which the follower kept a certain distance from the precedent. In the serial-follower condition, the location of the (*i* + 1)-th individual was sampled uniformly at random from the *surface* of the sphere with a radius of 1.5 centered at the i-th individual’s location, which ensured that the sampled location was bounded in the domain (by rejection sampling; Figure 6e). Likewise, the spoke individuals in the hub-spoke condition were located uniformly at random on the surface of the sphere with a radius of 1.5 centered at the hub individual (Figure 6f). In all the simulations, the location of the non-follower (e.g., the first individual in the serial condition and the hub individual in the hub-spoke condition) was sampled uniformly randomly from the domain.

For each of the simulations, 100,000 samples were generated, with each containing 3D coordinates of the five individuals.^4^ All samples were used for training and attention analysis. The researchers did not hold out a test portion of the data, as the purpose of the study was not to test the general applicability of the model predictions, but rather to reveal the dependency relations in the data (just as in linear regression tests).

In addition to the five-agent simulations resembling our real data from Japanese macaques, larger groups of 25 and 50 agents were studied using the proximal serial- and spoke-follower patterns (Figure 6c,6d; the same bounded space was used for the location domain). They were meant to test the scalability of the proposed method.

Note that in all the simulations, the location samples were the only information available for the neural network analyzer. Furthermore, the settings of the simulations, including the types of inter-individual relations and the form of the conditional distribution, were not hard-coded in the analyzer, unlike with other classical analyses that assume a limited hypothesis space including the correct generative model (e.g., Bayesian inference). For example, when analyzing the data from the hub-relation simulations, the network had to (i) discover the hub structure among many other possible relation patterns behind the data, and (ii) identify the hub individual while the five agents were always close to one another, without being tricked by different distributional patterns (proximal or distant).

Although the simulated data took on continuous values (residing in the bounded subspace of the 3D Euclidean space), it has been reported that neural networks are often better at making predictions of discrete categories (due to the flexibility of categorical distributions; van den Oord et al., 2016a,b, similar discrete predictions of location data were also adopted by Takeda and Komatani, 2016). Accordingly, the domain was split into 400 cubes (0.5 × 0.5 × 0.5), and the network was trained to make a discrete prediction of the cube containing the target’s location.

To assess the statistical significance of the estimated attention weights, they were compared with 100, 000 uniformly random weights sampled from Dirichlet(1,…, 1) (Ewens, 2003). Divergence of the estimated/random weights from the gold-standard weights was measured by the Kullback-Leibler (KL) divergence (Kullback and Leibler, 1951, see the supporting information S2 for the definition of the gold-standard weights):

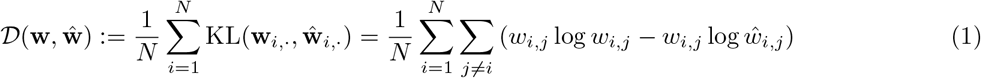

where *w_i,j_* and *ŵ_i,j_* denote the gold-standard and estimated/random attention weights respectively that were assigned to the referent individual *j* upon the prediction of the target individual *i*.

#### 2.2.2 Real Data

In addition to the simulated data described in §2.2.1, we analyzed real 3D location data of five captive Japanese macaques in the Primate Research Institute, Kyoto University (KUPRI), Japan. The subjects were two adult males and three adult females. Location recordings of the five macaques were conducted for two weeks in an outdoor group cage (4 × 5 × 2.5 m) where the subjects were allowed to freely move and interact with one another. Each subject carried two Bluetooth® Low-Energy beacons, and the 3D location was estimated based on the signals coming from these beacons (Quuppa Intelligent Locating System™). See Morita et al. (2020) for more details on the data collection procedures.

Although the system sampling frequency was set at 9 Hz, the actual rate was unstable due to uncontrollable signal interference and reflection perturbation. The sampling from the 2 × 5 = 10 beacons did not synchronize their timing. Therefore, we took the median value of the samples from each individual (i.e., from the two beacons) for each dimension collected at every 3000 msec interval. The medians over the same interval constituted the input-output pair for the neural network.^5^ We note that taking medians decreased time resolution of the collected data; however, temporal information was ignored in the analysis. The preprocessing steps above yielded a total of 327,592 data points.^6^ Just as in the simulations, all data were used for both the model training and attention analysis without holding out a test portion, and the target locations were discretized into 400 cubes (0.5 × 0. 5 × 0. 5 m).

## 3 Results

The attention-based analysis recovered the relational structures in the simulations (serial, hub-spoke, and other connective patterns) on a statistically significant level *(p* ≤ 0.00195) regardless of the distance pattern between the agents (proximal or distant). Given the data from the serial follower simulation, the analyzer correctly assigned heavier weights to the relations between the individuals with successive indices (e.g., “2” and “3”; Figure 7a,c, 8a,c). On another note, the hub-spoke simulation concentrated the attention weights to the hub individual when predicting the spoke dependents and evenly distributed the weights to the spokes when predicting the hub (Figure 7b,d, 8b,d). See the supporting information S1.2 for the results of the other relation patterns. Note that the analysis was non-parametric, as all the different simulations were analyzed using identical free parameters.

**Figure 7:**
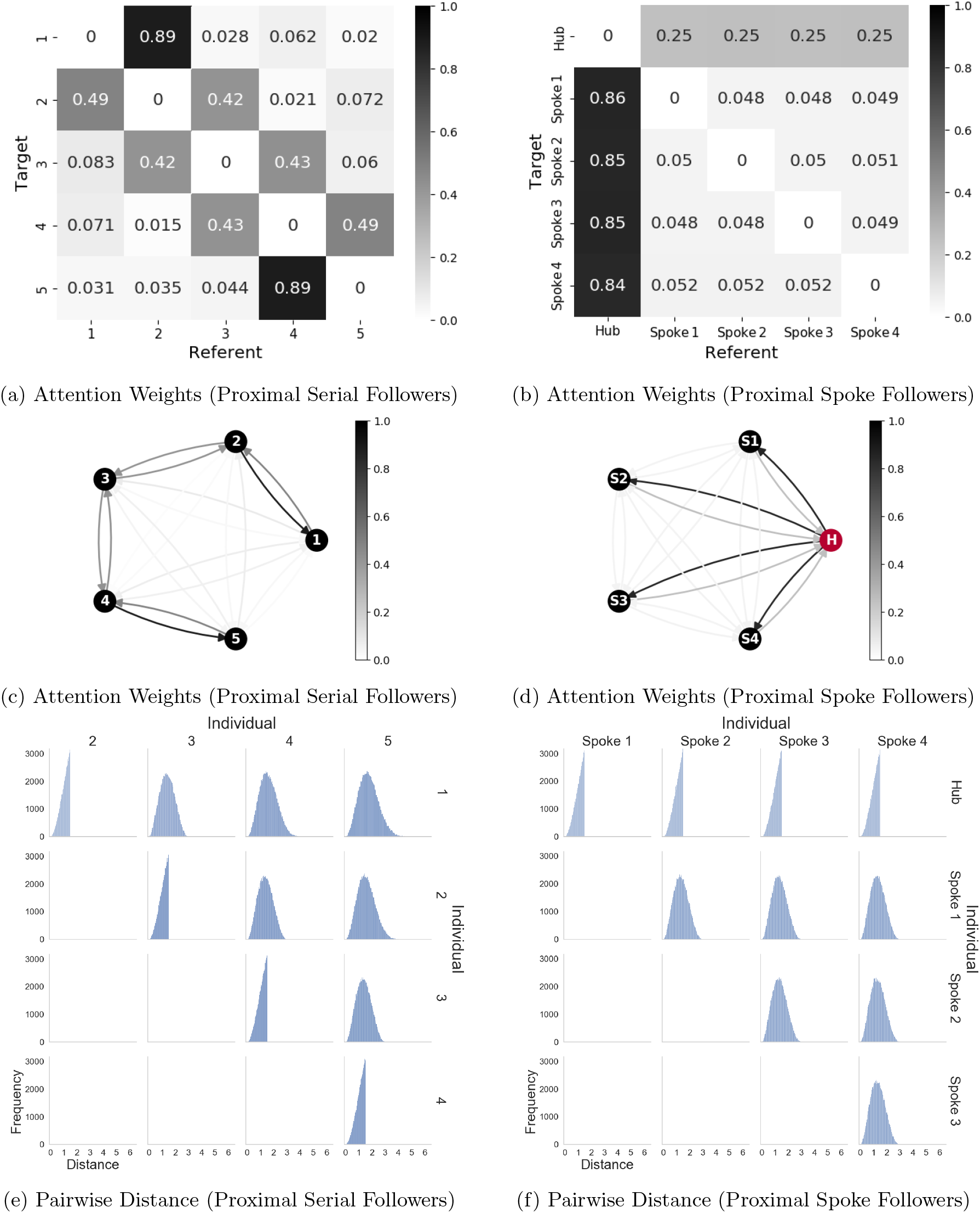
Analysis of proximal-follower simulations (left column: serial followers, Figure 6c; right column: spoke followers, Figure 6d). Top (a,b): Heatmap representation of average attention weights assigned to referent individuals (columns) upon the prediction of the location of target individuals (rows). Middle (c,d): Graphical representation of average attention weights assigned to referent individuals upon the prediction of the location of target individuals (i.e., the same data as the top, presented in a different format). Each arc runs from a referent to a target individual, with attention weight represented by color density. Bottom (e,f): Distribution of the pairwise 3D Euclidean distance between the individuals.

**Figure 8:**
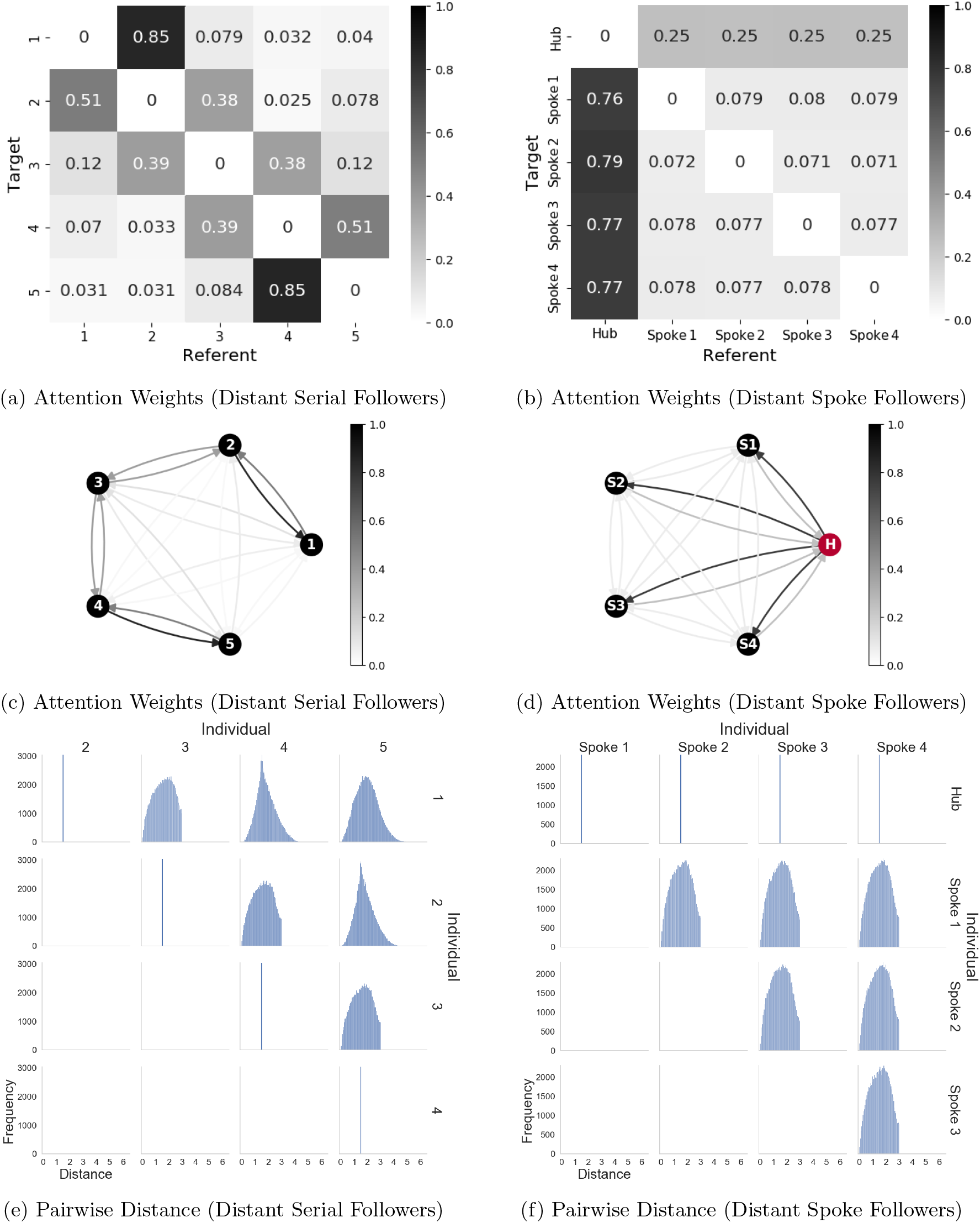
Analysis of distant-follower simulations (left column: serial followers, Figure 6e; right column: spoke followers, Figure 6f). Top (a,b): Heatmap representation of average attention weights assigned to referent individuals (columns) upon the prediction of the location of target individuals (rows). Middle (c,d): Graphical representation of average attention weights assigned to referent individuals upon the prediction of the location of target individuals (i.e., the same data as the top, presented in a different format). Each arc runs from a referent to a target individual, with attention weight represented by color density. Bottom (e,f): Distribution of the pairwise 3D Euclidean distance between the individuals.

Unlike our proposed method, traditional distance-based analyses cannot capture the correct relational patterns. Specifically, the relations among distant followers cannot be captured from their Euclidean distance, which was by definition constant between related individuals (at 1.5), while non-related individuals became closer for approximately 50% of the time (Figure 8e,f). Hence, non-related individuals would be incorrectly identified as related if, for example, the approaches under 1.5 were counted. Distance-based analyses would also be less robust for the proximal patterns. While the related individuals were at most 1.5 apart (by definition), the non-related individuals were located closer than 1.5 for 39-63% of the time (Figure 7e,7f). Thus, it is hard to achieve a robust contrast between the related and non-related individuals based on the distance information, especially to the extent achieved by the attention-based analysis (Figure 7a-d).

It should be noted that the detected relations are asymmetric but do not correspond to the causal structures in the simulations. Instead, the model computes the conditional probability of the location of each target individual given that of the referent individuals, and the attention weights represent the contribution of each referent individual to the computation of conditional probability. For example, the locations of the individual indexed “1” in the serial-follower simulations and the hub individual in the spoke-follower simulations were sampled unconditionally from the domain, without reference to any other individuals. However, the model can locate these individuals more accurately if it refers to their followers (the second and spoke individuals); in other words, the conditional probability, 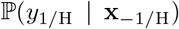 (where *y*_1/H_, is the (discretized) location of the leading/hub individual and **x**_-1/H_ is the (continuous-valued) location of the other individuals), rewritten as follows by Bayes’ Theorem:

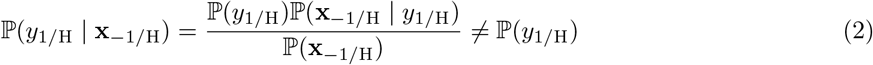

because 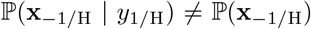 (i.e., the follower individuals are dependent on the leading/hub individual). Thus, non-zero attention weights should be assigned to the follower individuals.

While the proposed method was not able to detect the original causal relations, the proposed method distinguished direct vs. indirect statistical dependencies in the simulations. For example, individuals indexed “1” and “3” are only conditionally independent given the location of the individual indexed “2”; that is, if the second individual is not considered, the first and third individuals are located more accurately via a reference to the location of each other and, thus, are statistically dependent. This implies that naive pairwise evaluation of statistical dependence would fail to achieve a global structure consisting of direct dependencies. By contrast, the attention-based analysis aims to identify the direct dependencies from a number of possible relational structures.

One clear problem with the proposed method is its scalability; the model predictions can become less robust as more agents are included in the analysis. The estimated attention weights clearly captured the correct serial and hub-spoke relations when the number of simulated agents was scaled up to 25 (Figure 9a,9b). However, the difference between weights assigned to related and unrelated individuals was extremely small when the analysis was performed on 50 agents in the hub-spoke relation (Figure 9d). It is noteworthy, though, that the predicted attention weights are still statistically significant (p < 0.00001).

**Figure 9:**
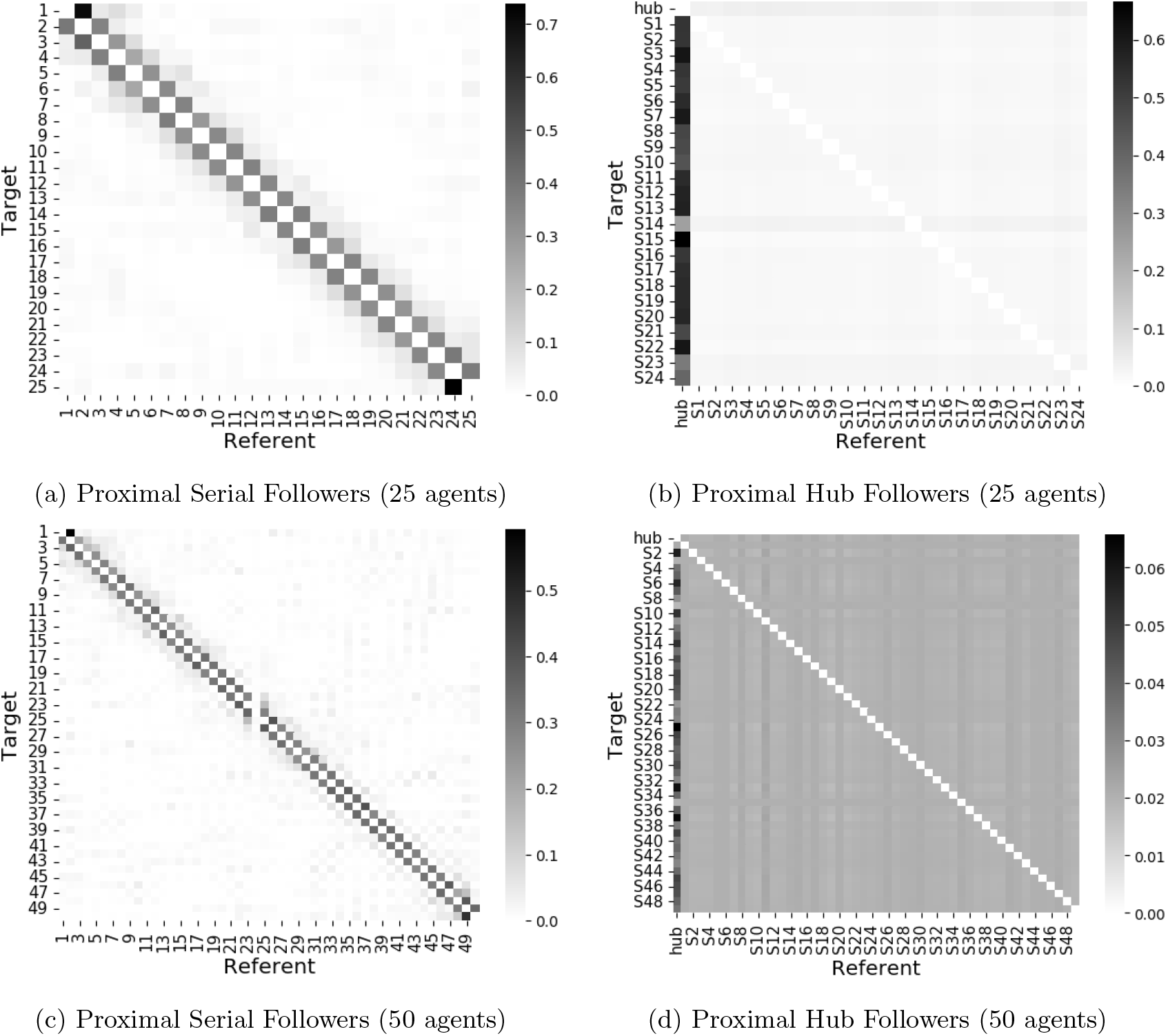
Average attention weights estimated from simulations with 25 and 50 agents (a,b and c,d respectively). The related individuals were positioned maximally 1.5 apart from one another (i.e., proximal relations) in all the simulations.

The attention-based analysis of the real data revealed that female macaques were more informative referents than males, especially when predicting the other females. The attention weight given to Male 1 as a referent was smaller than that of all the females, except when Male 2 was predicted (where Female 1 collected zero attention).^7^ Furthermore, Male 2 was almost never used as a referent (Figure 10a,b). It is noteworthy, however, that the males did not move independently of the other individuals. Predictions of the males’ locations were based on three other individuals (Male 1 was dependent on Female 1–3, and Male 2 was dependent on Female 2 and 3 and Male 1). While the neural network was unable to flag independence, as the attention weights were never all zeros, these detected dependencies were not redundant, because the network outperformed the no-referent baseline. As shown in Figure 10c, the average prediction loss (i.e., negative log probability) of the attention model was smaller than the baseline across all individuals.

**Figure 10:**
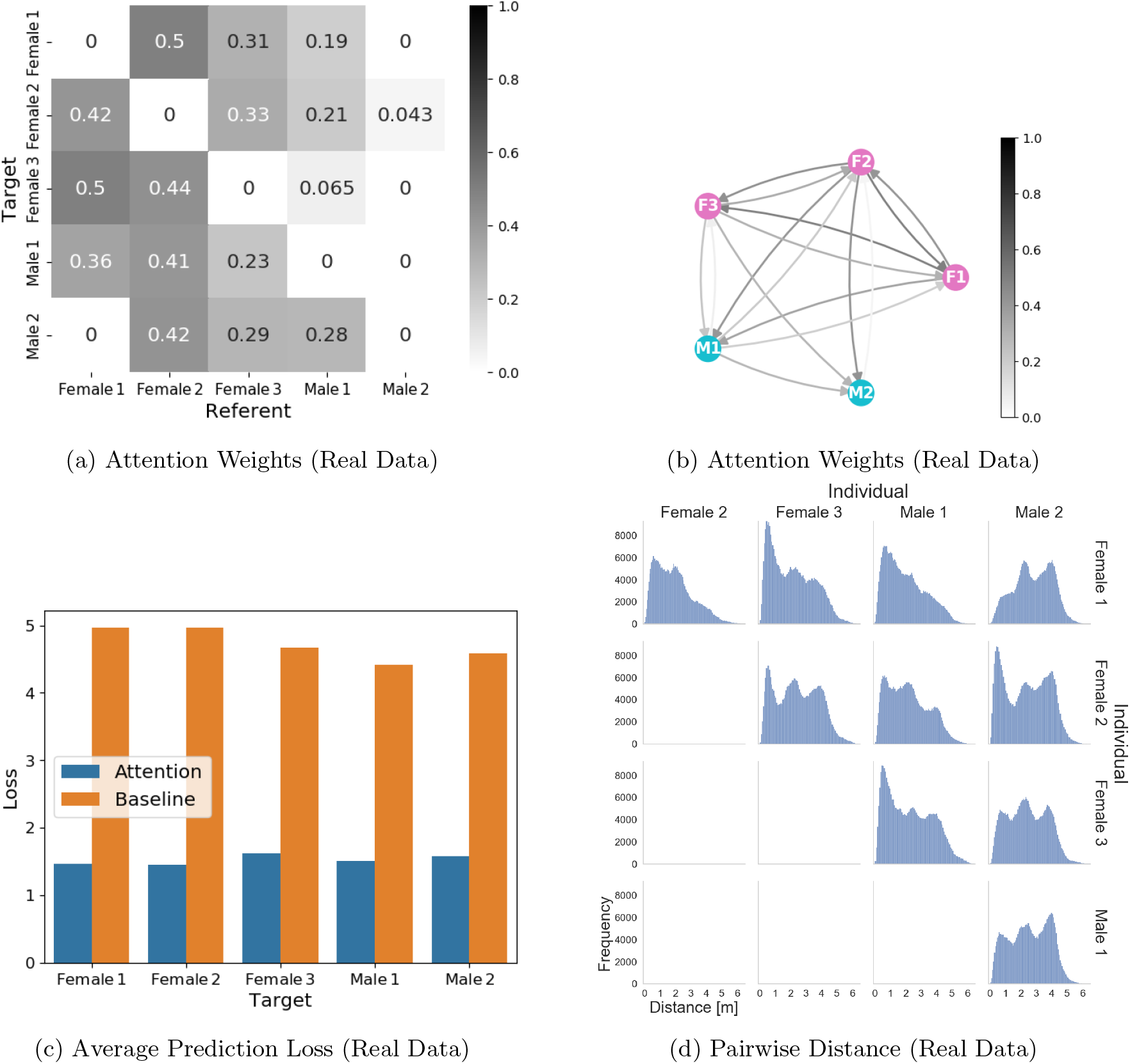
Analysis of real data collected from Japanese macaques. (a) Heatmap representation of average attention weights assigned to referent individuals (columns) upon the prediction of the location of target individuals (rows). (b) Graphical representation of the average attention weights assigned to referent individuals upon prediction of the location of target individuals (i.e., the same data as a, presented in a different format). Each arc runs from a referent to a target individual, with the attention weight represented by color density. (c) Average prediction loss (negative log probability) of the attention-based neural network (blue) and the no-referent baseline (orange). (d) Distribution of the pairwise 3D Euclidean distance between the individuals.

As with the simulation data, the real data challenge traditional distance-based analyses. The Euclidean distance between the macaques showed three frequency peaks: approximately 0.0–1.5, 1.5–3.0, and 3.0–4.5 m (Figure 10d). These multimodal distributions can originate from social relations among macaques or from other factors: for example, a small climbing frame and a hammock were installed around the center and peripherals of the experimental cage, and the second peak of the distance distribution could be caused by monkeys sitting on those facilities. In the current case, the canonical proximity analysis, which focused only on the leftmost peaks, seems to agree with the attention-based analysis. For example, both analyses suggest that Male 2 (taking greater distances from the others and receiving almost zero attention as a referent) was isolated from other macaques. Note, however, that the ignorance of the second and third peaks in the distributions was non-trivial and unjustified unless the more non-parametric analyses were performed.

## 4 Discussion

In this study, the attention analysis correctly identified the latent relational patterns behind the simulation, including serial, hub-spoke and other random relations, which was far beyond the chance level. It should be noted that the analyses were performed without prior assumptions about the possible types of inter-individual relations or random distributions involved. Compared with traditional distance/closeness-based analysis (Silk et al., 2006b,a; Croft et al., 2008; Clark, 2011; Boyland et al., 2013; Castles et al., 2014; Krause et al., 2015; Schofield et al., 2019; Gelardi et al., 2020), this method is likely more effective in visualizing latent dependency patterns, especially when little is known about the data properties. Specifically, analysis of the proximal co-appearance cannot identify related individuals when they keep their distance from one another, while attention-based analysis successfully detected the latent relations using the same values on the free parameters as used for the proximal pairwise relations. It is of note, however, that the goal of the attention-based analysis is not to *replace* conventional analyses, including that of geometric proximity; instead, the proposed method aims to act as a starting point in a series of data analyses, and measurement of the Euclidean distance and statistics of social activities (e.g., grooming) would still be informative and important for obtaining a detailed understanding of the detected relationships (Figure 1).

This study also demonstrated that the attention analysis is applicable to raw real data; in this instance, data from a case study of Japanese macaques was used for the analysis. We found that the location of females was more informative than that of males for locating the other female individuals. This result is consistent with the female-oriented social system of the macaques: Japanese macaques are a female-philopatric species (represented by parturition in the natal group and male emigration), and females usually build firmer and more stable social relationships with one another than do males (e.g., matrilineal social rank succession; Yamada, 1963; Furuichi, 1984; Mitani, 1986; Nakagawa et al., 2010). The females of the species are also reported to exhibit tighter and more frequent spatial cohesion (Otani et al., 2014), so knowing the location of some females will increase the ability to locate other females in the same group. In addition, our earlier research on the same data showed that the movements of males (i.e., trajectories of the 3D location) reflect a greater degree of individuality than do the movements of the females (Morita et al., 2020). Due to the idiosyncratic movements, the locations of males may not be informative predictors of other individuals’ locations. The analysis of the real data also revealed that one individual (Male 2) was almost never attended by the others. The discovery of such distinguished individuals is useful for building new hypotheses about animal societies and planing behavioral experiments targeting them. The attention-based analysis detects (non-)related individuals without presupposing specific types of relations, so the detected relations may reveal hitherto unknown patterns of social structures that are not necessarily based on proximal relations, in combination with other analytical tools (Figure 1).

Attention-based neural networks are now being adopted in biological studies. Yuan and Jia (2019) used the attention mechanism to implement a seizure detector based on electroencephalogram (EEG) data. They showed that the attention mechanism can locate EEG channels that specifically diagnose seizures. Similarly, Maekawa et al. (2020) analyzed movements of worms and mice using the attention mechanism and highlighted their characteristic movements under different experimental conditions. These studies reemphasize that attention-based data analyses target situations in which researchers do not know which aspects/features/portions of the data are most important; this limitation is overcome by implementing a data-driven detection of relevant features. Although such non-parametric analyses have been attempted using information-theoretical methods, those approaches often have limitations compared to deep learning. For example, Chen et al. (2019, 2020) inferred the causal relationship among pigeons from GPS data based on causation entropy (i.e., mutual information between an individual at a discrete time *t* and another at time *t* +1, conditioned on some other individuals at t Sun et al., 2015). While the information-theoretical analysis is non-parametric in principle, empirical computation includes an estimation of probability distributions and, therefore, can spoil the non-parametricity. Chen et al. (2019) defined probability distributions based on a 2D Euclidean proximity metric, which would miss distant dependencies that are similar to the simulations performed in this study. Chen et al. (2020) instead used 2D velocity directions that were discretized into eight regions evenly divided in the polar coordinate (i.e., 0-45°, 45-90°, …, and 315-360°). This discretization was sparse compared with that of the present study (8 vs. 400 levels), because fine-grained discretization, specifically of the conditioning variables representing the referent individuals, makes traditional statistical estimation difficult to perform. By contrast, the attention-based analysis, which utilized neural networks, can exploit continuous-valued data for the conditioning variables (raw data or embeddings of discrete data), and sparse discretization is unnecessary. Therefore, attention-based analysis can be more detailed and non-parametric than traditional information-theoretical approaches.

Attention analysis does not exploit any behavioral patterns that are particular to Japanese macaques, and researchers can use it for location data of other animals. The analysis is particularly appropriate when the use of conventional distance/closeness metrics is unsupported. Geometric proximity can be a reasonable proxy for observations of some social behaviors, such as grooming (Gelardi et al., 2020), but species without such social behaviors may exhibit a relationship at a distance. We further emphasize that attention analysis is not limited to location data; the flexibility of the neural network allows for its application to a wide variety of data with interacting components. For example, vocal communication has been of major interest to behavioral biologists and evolutionary linguists (Levinson, 2016). Attention analysis could potentially reveal latent communication flow when it is applied to vocalization recordings of multiple individuals. However, the proposed method needs to be extended and refined for broader applications and for more detailed analyses of location/movement. It was shown that model predictions can be less robust and harder to interpret as a greater number of individuals are included in the analysis. The proposed method would be effective for behavioral data including 10-25 individuals (e.g., Nagy et al., 2010; Strandburg-Peshkin et al., 2015), but limited to a larger group with hundreds of agents, such as social insects. Robust inference via the attention mechanism is an active research area (Martins and Astudillo, 2016; Niculae and Blondel, 2017; Laha et al., 2018; Niculae et al., 2018; Peters et al., 2018), and such updates should be incorporated into biological applications. Besides the scalability issue, the proposed method does not process temporal changes in location, whereas temporal analysis is a crucial part of analyzing dynamic behaviors and vocal communication. Since the attention mechanism was developed for NLP, however, processing the time series data should not be difficult (see Vaswani et al., 2017; Shaw et al., 2018; Dai et al., 2019, for popular ways to encode time information).

## Acknowledgements

We appreciate the animal care given by our technicians and research assistants as well as their suggestions and support for this project. In particular, we wish to express special thanks to Norihiko Maeda, Mayumi Morimoto, and Takayoshi Natsume for the animal arrangements, and Panasonic Solution Technologies Co. Ltd. for technical support with the equipment. We also acknowledge the useful comments and suggestions about the analysis from Zin Arai and Hiroshi Takeuchi. This study was performed under the Cooperative Research Program at KUPRI (2018-C-27, 2019-B-27, 2020-B-17). This study was mainly funded by the Japan Science and Technology Agency, Core Research for Evolutional Science and Technology 17941861 (#JPMJCR17A4) and partly by MEXT Grant-in-aid for Scientific Research on Innovative Areas #4903 (Evolinguistic), 17H06380.

## Ethical Statement

All procedures were reviewed and approved by the Animal Welfare and Care Committee of KUPRI (Permission # 2018-203) and complied with the institutional guidelines (Primate Research Institute, 2010).

## Author contributions

Project organization: IA IM HK; Animal arrangements: HK SA NSH; Apparatus building: AT IM HK; Data acquisition: AT IM HK; Animal cares: AK AT IM HK; Data management; HK AT TM; Computational modeling: TM; Manuscript writing: IA IM HK TM.

## Data Availability

The data and code used in this study will become available in Mendelay Data and GitHub respectively when it is accepted for publication.

## Footnotes

1 As reported in §2.2, the location of the target individual was discretized to improve the performance of the neural networks (van den Oord et al., 2016a,b) and simplify the maximum likelihood training of the baseline model (i.e., the occurrences of each discretized location value were counted and the gathered data were normalized).

2 The scaling of the dot product of the key and query is based on the hidden dimensionality of each head. That is, the scalar is 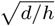 where *d* is the hidden dimensionality before head-splitting and *h* is the number of heads.

3 Note that **E**^(*p*,1)^, post-attention embeddings in the first attention layer, duplicate the role of **E**^(*q*,2)^, which constitutes the query input of the second layer. One of them can be removed, in principle.

4 The number of different target-referents pairs observed by the neural network was 100, 000*N* (where *N* ∈ {5, 25, 50} stands for the number of individuals), since all possible partitions of the *N* individuals were treated as one target and N – 1 referents in the analysis.

5 The network simultaneously analyzed all possible choices for one individual as the prediction target and for four individuals as the referents.

6 Again, the researchers analyzed all possible partitions of the five individuals into one target and four referents; the number of different target-referent pairs observed by the neural network was therefore 5 × 327, 592 = 1, 637, 960.

7 The average attention weighted to Male 1, upon the predictions of Male 2, appeared to be equal to the attention weighted to Female 3, but was statistically significantly smaller given the 10,000 samples of bootstrapping.

## References

Bahdanau, D., Cho, K., and Bengio, Y. (2015). Neural machine translation by jointly learning to align and translate. In 3rd International Conference on Learning Representations, ICLR 2015, San Diego, CA, USA, May 7-9, 2015, Conference Track Proceedings.

Boyland, N. K., James, R., Mlynski, D. T., Madden, J. R., and Croft, D. P. (2013). Spatial proximity loggers for recording animal social networks: consequences of inter-logger variation in performance. Behavioral Ecology and Sociobiology, 67(11):1877–1890.

Canteloup, C., Puga-Gonzalez, I., Sueur, C., and van de Waal, E. (2020). The effects of data collection and observation methods on uncertainty of social networks in wild primates. American Journal of Primatology, 82(7):e23137.

Castles, M., Heinsohn, R., Marshall, H. H., Lee, A. E., Cowlishaw, G., and Carter, A. J. (2014). Social networks created with different techniques are not comparable. Animal Behaviour, 96:59–67.

Chen, D., Kang, M., and Yu, W. (2020). Probabilistic causal inference for coordinated movement of pigeon flocks. EPL (Europhysics Letters), 130(2):28004.

Chen, D., Wang, Y., Wu, G., Kang, M., Sun, Y., and Yu, W. (2019). Inferring causal relationship in coordinated flight of pigeon flocks. Chaos: An Interdisciplinary Journal of Nonlinear Science, 29(11):113118.

Chepko-Sade, B., Reitz, K. P., and Sade, D. S. (1989). Sociometrics of macaca mulatta iv: Network analysis of social structure of a pre-fission group. Social Networks, 11(3):293–314. Special Issue on Non-Human Primate Networks.

Clark, F. E. (2011). Space to choose: network analysis of social preferences in a captive chimpanzee community, and implications for management. American Journal of Primatology, 73(8):748–757.

Croft, D. P., James, R., and Krause, J. (2008). Exploring Animal S’ocial Networks. Princeton University Press.

Cybenko, G. (1989). Approximation by superpositions of a sigmoidal function. Mathematics of Control, Signals and Systems, 2(4):303–314.

Dai, Z., Yang, Z., Yang, Y., Carbonell, J., Le, Q., and Salakhutdinov, R. (2019). Transformer-XL: Attentive language models beyond a fixed-length context. In Proceedings of the 57th Annual Meeting of the Association for Computational Linguistics, pages 2978–2988, Florence, Italy. Association for Computational Linguistics.

Devlin, J., Chang, M.-W., Lee, K., and Toutanova, K. (2018). BERT: Pre-training of deep bidirectional transformers for language understanding. arXiv:1810.04805.

Dore, K. M., Hansen, M. F., Klegarth, A. R., Fichtel, C., Koch, F., Springer, A., Kappeler, P., Parga, J. A., Humle, T., Colin, C., Raballand, E., Huang, Z.-P., Qi, X.-G., Di Fiore, A., Link, A., Stevenson, P. R., Stark, D. J., Tan, N., Gallagher, C. A., Anderson, C. J., Campbell, C. J., Kenyon, M., Pebsworth, P., Sprague, D., Jones-Engel, L., and Fuentes, A. (2020). Review of GPS collar deployments and performance on nonhuman primates. Primates.

Ewens, W. J. (2003). On estimating p values by the Monte Carlo method. American journal of human genetics, 72(2):496–498.

Farine, D. R. and Whitehead, H. (2015). Constructing, conducting and interpreting animal social network analysis. Journal of Animal Ecology, 84(5):1144–1163.

Fehlmann, G. and King, A. J. (2016). Bio-logging. Current Biology, 26(18):R830–R831.

Furuichi, T. (1984). Symmetrical patterns in non-agonistic social interactions found in unprovisioned Japanese macaques. Journal of Ethology, 2(2):109–119.

Gelardi, V., Godard, J., Paleressompoulle, D., Claidiere, N., and Barrat, A. (2020). Measuring social networks in primates: wearable sensors versus direct observations. Proceedings of the Royal Society A: Mathematical, Physical and Engineering Sciences, 476(2236):20190737.

Haddadi, H., King, A. J., Wills, A. P., Fay, D., Lowe, J., Morton, A. J., Hailes, S., and Wilson, A. M. (2011). Determining association networks in social animals: choosing spatial-temporal criteria and sampling rates. Behavioral Ecology and Sociobiology, 65(8):1659–1668.

Heupel, M. R., Semmens, J. M., and Hobday, A. J. (2006). Automated acoustic tracking of aquatic animals: scales, design and deployment of listening station arrays. Marine and Freshwater Research, 57(1):1–13.

Hornik, K. (1991). Approximation capabilities of multilayer feedforward networks. Neural Networks, 4(2):251–257.

Jin, L., Gupta, M. M., and Nikiforuk, P. N. (1995). Universal approximation using dynamic recurrent neural networks: discrete-time version. In Proceedings of ICNN’95 - International Conference on Neural Networks, volume 1, pages 403–408.

King, A. J., Clark, F. E., and Cowlishaw, G. (2011a). The dining etiquette of desert baboons: the roles of social bonds, kinship, and dominance in co-feeding networks. American Journal of Primatology, 73(8):768–774.

King, A. J., Fehlmann, G., Biro, D., Ward, A. J., and Furtbauer, I. (2018). Re-wilding collective behaviour: An ecological perspective. Trends in Ecology & Evolution, 33(5):347–357.

King, A. J., Sueur, C., Huchard, E., and Cowlishaw, G. (2011b). A rule-of-thumb based on social affiliation explains collective movements in desert baboons. Animal Behaviour, 82(6):1337–1345.

Krause, J., James, R., Franks, D. W., and Croft, D. P., editors (2015). Animal Social Networks. Oxford University Press.

Kullback, S. and Leibler, R. A. (1951). On information and sufficiency. The Annals of Mathematical Statistics, 22(1):79–86.

Laha, A., Chemmengath, S. A., Agrawal, P., Khapra, M. M., Sankaranarayanan, K., and Ramaswamy, H. G. (2018). On controllable sparse alternatives to softmax. In Proceedings of the 32Nd International Conference on Neural Information Processing S’ystems, NIPS’18, pages 6423–6433, USA. Curran Associates Inc.

Levinson, S. C. (2016). Turn-taking in human communication - origins and implications for language processing. Trends in Cognitive Sciences, 20(1):6–14.

Maekawa, T., Ohara, K., Zhang, Y., Fukutomi, M., Matsumoto, S., Matsumura, K., Shidara, H., Yamazaki, S. J., Fujisawa, R., Ide, K., Nagaya, N., Yamazaki, K., Koike, S., Miyatake, T., Kimura, K. D., Ogawa, H., Takahashi, S., and Yoda, K. (2020). Deep learning-assisted comparative analysis of animal trajectories with deephl. Nature Communications, 11(1):5316.

Martins, A. and Astudillo, R. (2016). From softmax to sparsemax: A sparse model of attention and multilabel classification. In Balcan, M. F. and Weinberger, K. Q., editors, Proceedings of The 33rd International Conference on Machine Learning, volume 48 of Proceedings of Machine Learning Research, pages 1614–1623, New York, New York, USA. PMLR.

Mitani, M. (1986). Voiceprint identification and its application to sociological studies of wild Japanese monkeys (*Macaca fuscata yakui*). Primates, 27(4):397–412.

Morita, T., Toyoda, A., Aisu, S., Kaneko, A., Suda-Hashimoto, N., Matsuda, I., and Koda, H. (2020). Animals exhibit consistent individual differences in their movement: A case study on location trajectories of Japanese macaques. Ecological Informatics, 56:101057.

Nagy, M., Akos, Z., Biro, D., and Vicsek, T. (2010). Hierarchical group dynamics in pigeon flocks. Nature, 464(7290):890–893.

Nakagawa, N., Nakamichi, M., and Sugiura, H. (2010). The Japanese Macaques. Primatology Monographs. Springer Japan.

Niculae, V. and Blondel, M. (2017). A regularized framework for sparse and structured neural attention. In Guyon, I., Luxburg, U. V., Bengio, S., Wallach, H., Fergus, R., Vishwanathan, S., and Garnett, R., editors, Advances in Neural Information Processing Systems 30, pages 3338–3348. Curran Associates, Inc.

Niculae, V., Martins, A., Blondel, M., and Cardie, C. (2018). SparseMAP: Differentiable sparse structured inference. In Dy, J. and Krause, A., editors, Proceedings of the 35th International Conference on Machine Learning, volume 80 of Proceedings of Machine Learning Research, pages 3799–3808, Stockholmsmässan, Stockholm Sweden. PMLR.

Otani, Y., Sawada, A., and Hanya, G. (2014). Short-term separation from groups by male Japanese macaques: Costs and benefits in feeding behavior and social interaction. American Journal of Primatology, 76(4):374–384.

Pays, O., Benhamou, S., Helder, R., and Gerard, J.-F. (2007). The dynamics of group formation in large mammalian herbivores: an analysis in the european roe deer. Animal Behaviour, 74(5):1429–1441.

Pérez, J., Marinkovic, J., and Barceló, P. (2019). On the turing completeness of modern neural network architectures. In 7th International Conference on Learning Representations, ICLR 2019, New Orleans, LA, USA, May 6-9, 2019.

Peters, B., Niculae, V., and Martins, A. F. T. (2018). Interpretable structure induction via sparse attention. In Proceedings of the 2018 EMNLP Workshop BlackboxNLP: Analyzing and Interpreting Neural Networks for NLP, pages 365–367, Brussels, Belgium. Association for Computational Linguistics.

Primate Research Institute (2010). The Care and Use of Laboratory Primates. The Primate Research Institute, Kyoto University, third edition.

Rutz, C., Burns, Z. T., James, R., Ismar, S. M. H., Burt, J., Otis, B., Bowen, J., and St Clair, J. J. H. (2012). Automated mapping of social networks in wild birds. Current Biology, 22(17):R669–R671.

Rutz, C. and Hays, G. C. (2009). New frontiers in biologging science. Biology Letters, 5(3):289–292.

Schofield, D., Nagrani, A., Zisserman, A., Hayashi, M., Matsuzawa, T., Biro, D., and Carvalho, S. (2019). Chimpanzee face recognition from videos in the wild using deep learning. Science Advances, 5(9).

Shaw, P., Uszkoreit, J., and Vaswani, A. (2018). Self-attention with relative position representations. In Proceedings of the 2018 Conference of the North American Chapter of the Association for Computational Linguistics: Human Language Technologies, Volume 2 (Short Papers), pages 464–468, New Orleans, Louisiana. Association for Computational Linguistics.

Silk, J., Alberts, S., and Altmann, J. (2006a). Social relationships among adult female baboons (*Papio cynocephalus*) II. variation in the quality and stability of social bonds. Behavioral Ecology and Sociobiology, 61:197–204.

Silk, J., Altmann, J., and Alberts, S. (2006b). Social relationships among adult female baboons (*Papio cynocephalus*) I. variation in the strength of social bonds. Behavioral Ecology and Sociobiology, 61:183–195.

Sosa, S., Sueur, C., and Puga-Gonzalez, I. (2020). Network measures in animal social network analysis: Their strengths, limits, interpretations and uses. Methods in Ecology and Evolution.

Strandburg-Peshkin, A., Farine, D. R., Couzin, I. D., and Crofoot, M. C. (2015). Shared decision-making drives collective movement in wild baboons. Science, 348(6241):1358–1361.

Sun, J., Taylor, D., and Bollt, E. M. (2015). Causal network inference by optimal causation entropy. SIAM Journal on Applied Dynamical Systems, 14(1):73–106.

Takeda, R. and Komatani, K. (2016). Sound source localization based on deep neural networks with directional activate function exploiting phase information. In 2016 IEEE International Conference on Acoustics, Speech and Signal Processing (ICASSP), pages 405–409.

van den Oord, A., Dieleman, S., Zen, H., Simonyan, K., Vinyals, O., Graves, A., Kalchbrenner, N., Senior, A., and Kavukcuoglu, K. (2016a). Wavenet: A generative model for raw audio.

van den Oord, A., Kalchbrenner, N., Espeholt, L., kavukcuoglu, k., Vinyals, O., and Graves, A. (2016b). Conditional image generation with PixelCNN decoders. In Lee, D. D., Sugiyama, M., Luxburg, U. V., Guyon, I., and Garnett, R., editors, Advances in Neural Information Processing Systems 29, pages 4790–4798. Curran Associates, Inc.

Vaswani, A., Shazeer, N., Parmar, N., Uszkoreit, J., Jones, L., Gomez, A. N., Kaiser, L. u., and Polosukhin, I. (2017). Attention is all you need. In Guyon, I., Luxburg, U. V., Bengio, S., Wallach, H., Fergus, R., Vishwanathan, S., and Garnett, R., editors, Advances in Neural Information Processing Systems 30, pages 5998–6008. Curran Associates, Inc.

Yamada, M. (1963). A study of blood-relationship in the natural society of the Japanese macaque. Primates, 4(3):43–65.

Yuan, Y. and Jia, K. (2019). FusionAtt: Deep fusional attention networks for multi-channel biomedical signals. Sensors, 19(11):2429.

