## supporting information for "Non-Parametric Analysis of Inter-Individual Relations Using an Attention-Based Neural Network"

### S1 Details on Random Relation Structures

This section describes the detailed procedure for sampling random relation structures from which simulated data are generated (S1.1), and also reports the results of the proposed analysis on the data in detail (S1.2).

#### S1.1 Data Sampling

Causal relation structures can be represented as directed acyclic graphs. To cover a wider variety of graphs than the manually constructed patterns (serial and spoke followers), we exhaustively studied nine non-isolated graphs whose maximal indegree was one (Figure S1.1; isomorphic patterns were not repeated).

We also experimented with ten random graphs with a greater maximal indegree. Specifically, the directed edge from $j$-th node to $i$-th node ($j < i$) was created with a probability of 0.5. Note that this procedure does not guarantee that the maximal indegree of the sampled graphs was at least two (i.e., it can repeat the serial, hub-spoke, and other patterns in which each agent followed at most one precedent). Accordingly, we filtered out graphs with a smaller indegree by rejection sampling. Similarly, sampled graphs were rejected when they were isomorphic to the previous samples or the serial/hub-spoke patterns that were manually selected. Finally, we also rejected sampled graphs when some of the nodes were isolated (i.e., no incoming or outgoing edge). The sampled graphs are reported in S1.2.

The graphs in Figure S1.1 and S1.2 were then used to generate location data of the individuals corresponding to the nodes. As described in the main text (§2.2.1), the location of each individual was uniformly randomly sampled from either the ball (proximal) or the sphere (distant) centered at the location of the individual corresponding to the parent node. Note that some of the nodes in the random graphs have multiple parents (Figure S1.2a). In such cases, one of the parent nodes was randomly selected and the child's location was sampled conditioned on that node.

#### S1.2 Results

Figure S1.3–S1.12 reports the results of the proposed analysis on each of the random graphs. The left (a,c,e) and right (b,d,f) columns report the same results in different formats (graph and heatmap respectively). The top rows (a,b) show the gold-standard weights defined in S2. The middle rows (c,d) report the average attention weights predicted from the proximal-follower simulations in which each agent was located within the ball centered at one of the conditioning agents. The bottom rows (e,f) show the weights predicted from the distant-follower simulations in which each agent kept a constant distance from the conditioning agents.

When the maximal indegree of the generative graphs was one (Figure S1.3–S1.11), the proposed analysis assigned greater attention weights to the directly related individuals than to the unrelated/non-directly related ones. When the graphs had a greater indegree (Figure S1.12–S1.21), on the other hand, some unrelated/non-directly related individuals received greater attention weights than those of the directly related pairs, and the KL divergence between the gold-standard and estimated attention weights increased ($0.1306 \pm 0.0361 \rightarrow 0.1844 \pm 0.0409$ for the proximal relations and $0.1716 \pm 0.0567 \rightarrow 0.2229 \pm 0.0407$ for the distant relations on average $\pm$ standard deviation). Nevertheless, the KL divergence was statistically significantly small in all simulations ($p \leq 0.00195$).

### S2 Gold-Standard Attention Weights

The statistical significance of estimated attention weights was assessed in reference to gold-standard weights. The gold-standard weights of each relation structure were optimally distributed over the referent individuals; given a directed acyclic connected graph corresponding to the generative model of the location data, the gold-standard weight on each referent node upon the prediction of each target node is proportional to the probability of using the edge between the two nodes. Consider the directed graph in Figure S2.1, for example, setting the node X as the target. It has three parents (A, B, and C) and they have the equal probability $\frac{1}{3}$ of conditioning X. The target node X also has two children, D and E. X is the only parent node of D, so it is used as a conditioner with probability 1. By contrast, E has another parent (F), and thus, the edge between

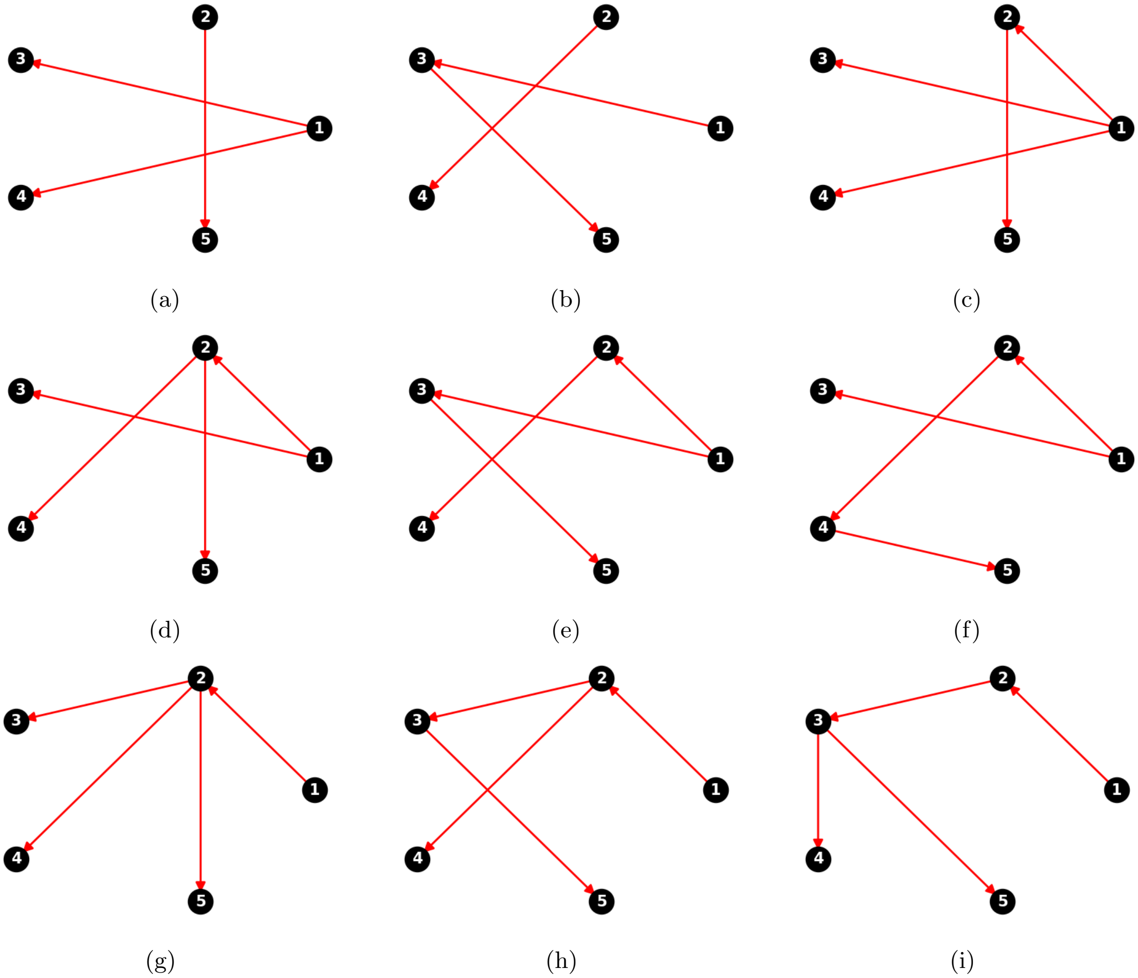


Figure S1.1: Directed acyclic graphs used for generating location data. The maximal indegree is one.

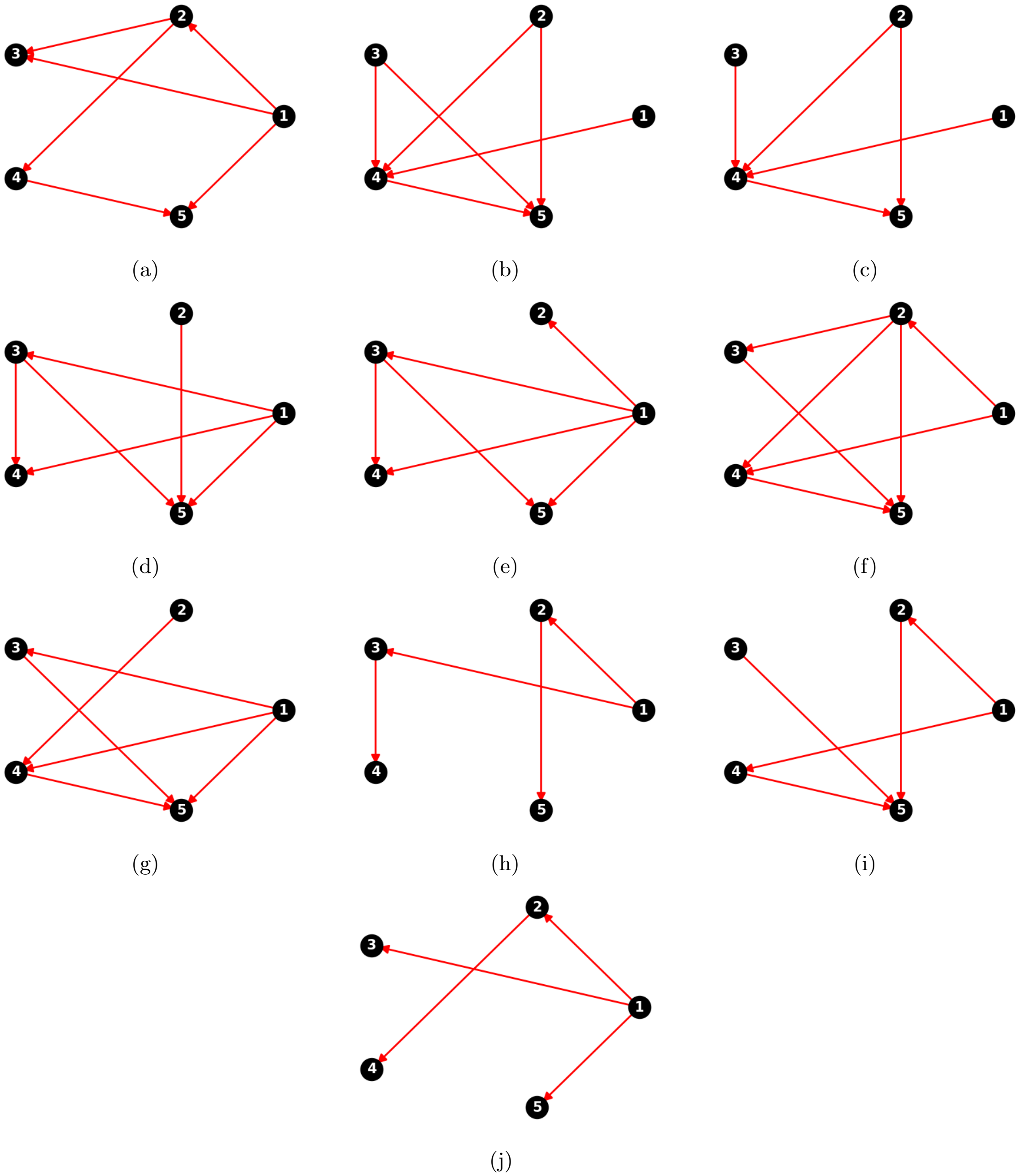


Figure S1.2: Random directed acyclic graphs used for generating location data. The maximal indegree is two or greater.

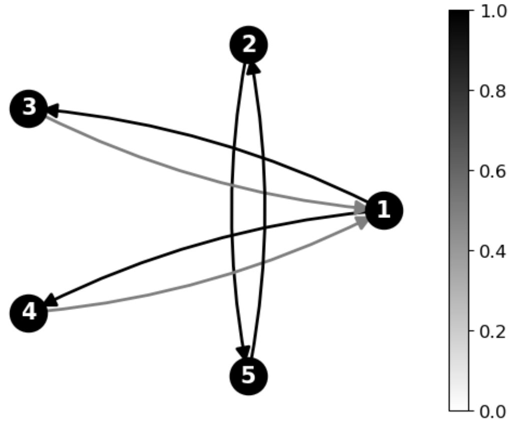


(a) Gold-Standard Weights

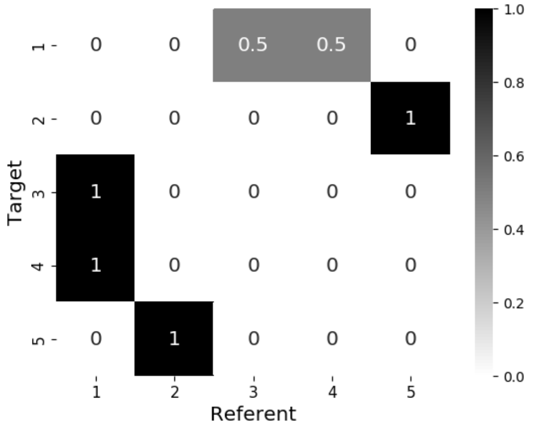


(b) Gold-Standard Weights

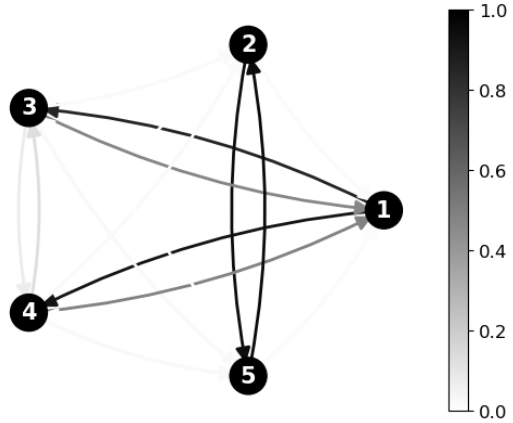


(c) Proximal Followers

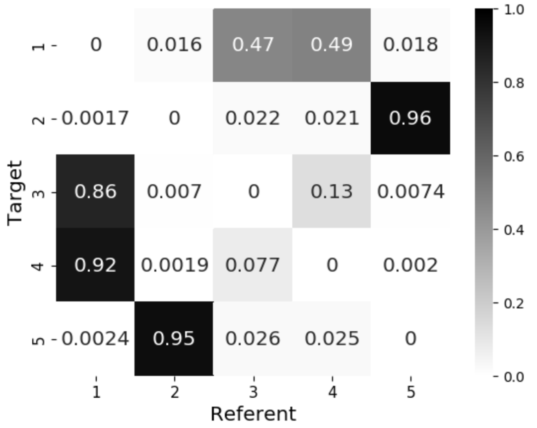


(d) Proximal Followers

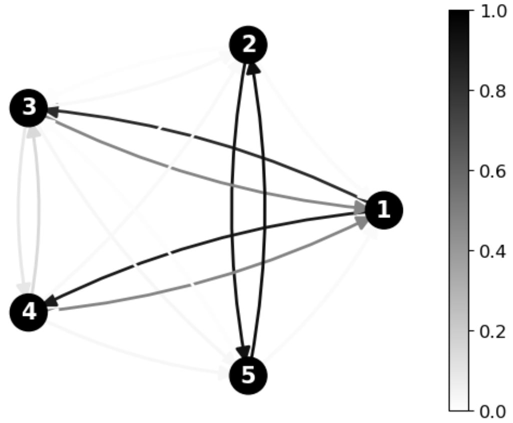


(e) Distant Followers

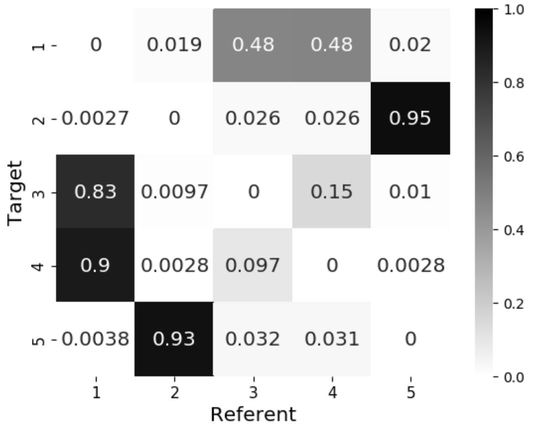


(f) Distant Followers

Figure S1.3: Gold-standard and average predicted attention weights for the random graph in Figure S1.1a. The KL divergence between gold-standard and predicted attention weights was 0.0751 for proximal followers ($p < 0.00001$) and 0.0919 for distant followers ($p < 0.00001$).

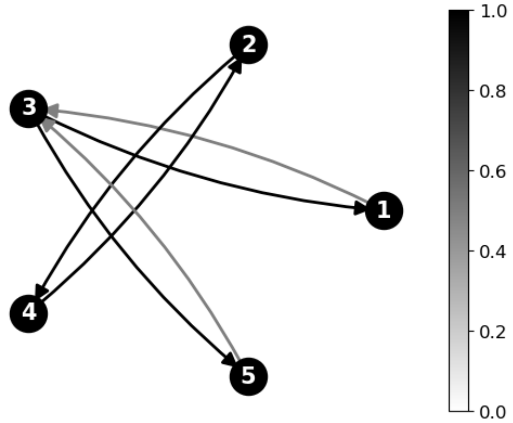


(a) Gold-Standard Weights

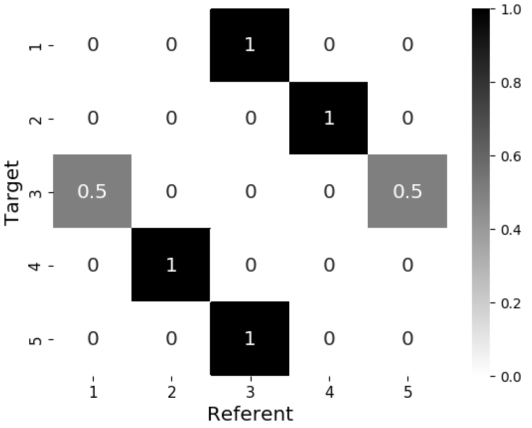


(b) Gold-Standard Weights

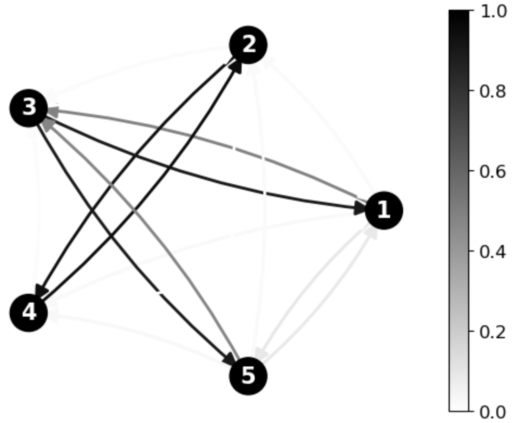


(c) Proximal Followers

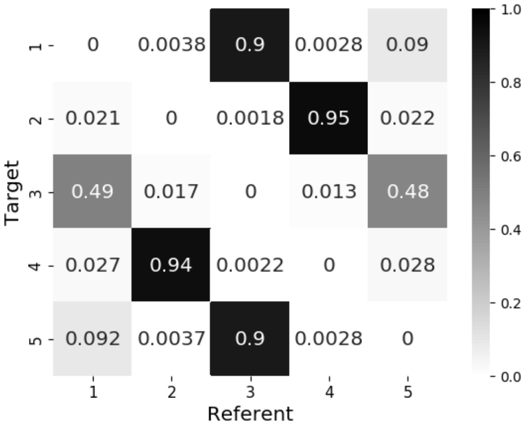


(d) Proximal Followers

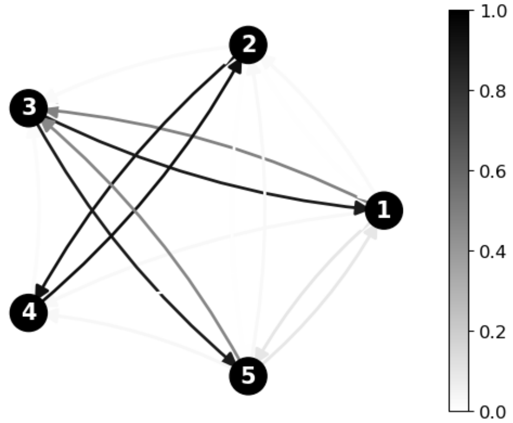


(e) Distant Followers

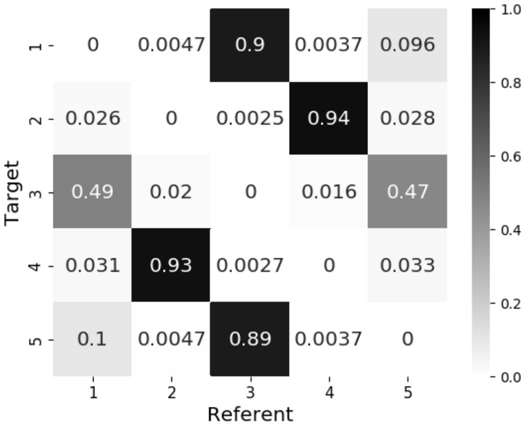


(f) Distant Followers

Figure S1.4: Gold-standard and average predicted attention weights for the random graph in Figure S1.1b. The KL divergence between gold-standard and predicted attention weights was 0.0681 for proximal followers ($p < 0.00001$) and 0.0779 for distant followers ($p < 0.00001$).

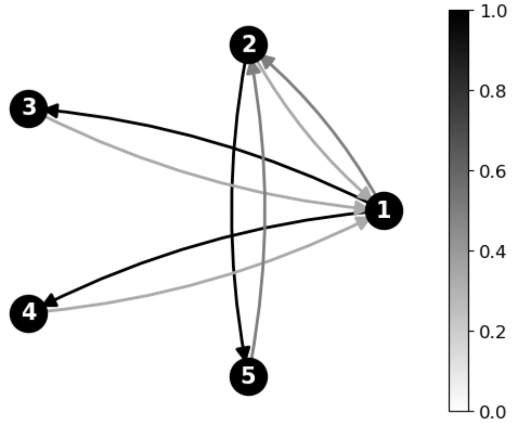


(a) Gold-Standard Weights

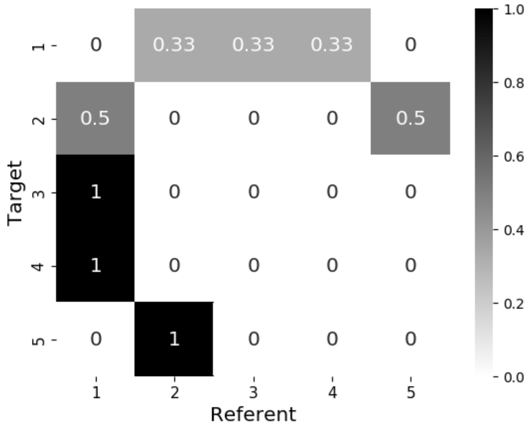


(b) Gold-Standard Weights

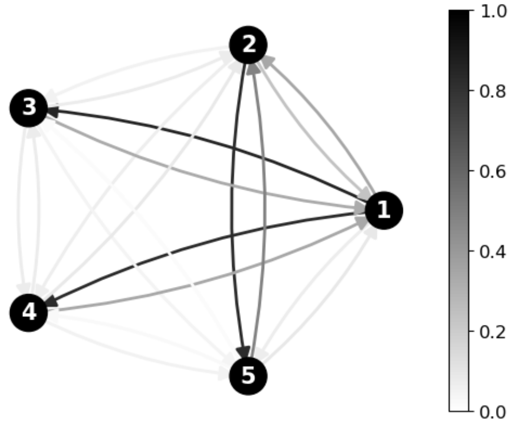


(c) Proximal Followers

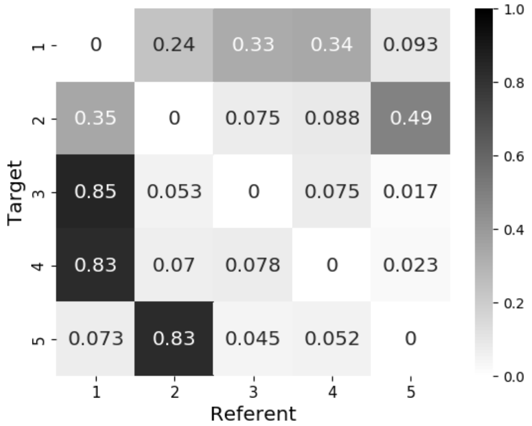


(d) Proximal Followers

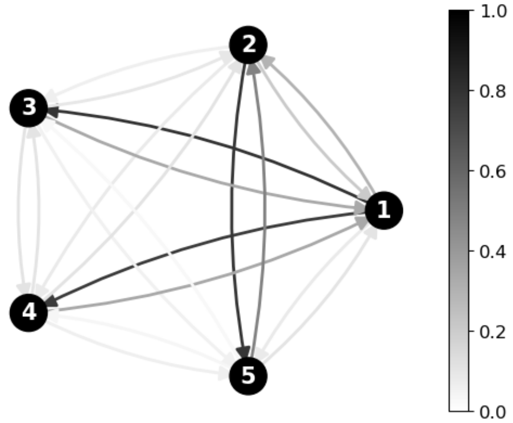


(e) Distant Followers

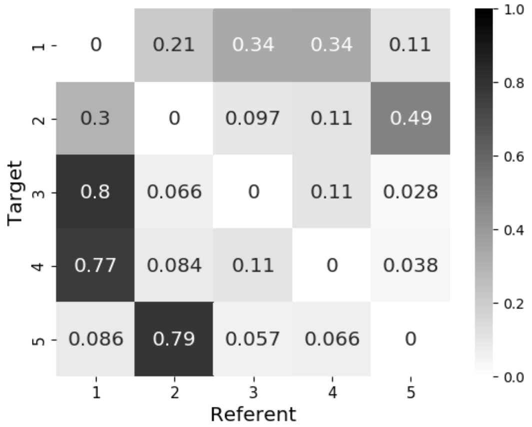


(f) Distant Followers

Figure S1.5: Gold-standard and average predicted attention weights for the random graph in Figure S1.1c. The KL divergence between gold-standard and predicted attention weights was 0.1668 for proximal followers ($p < 0.00001$) and 0.2264 for distant followers ($p = 0.00003$).

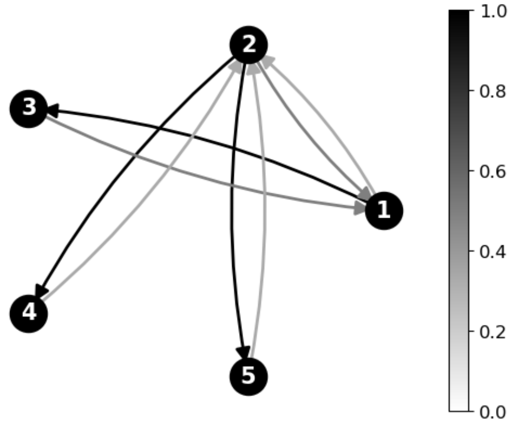


(a) Gold-Standard Weights

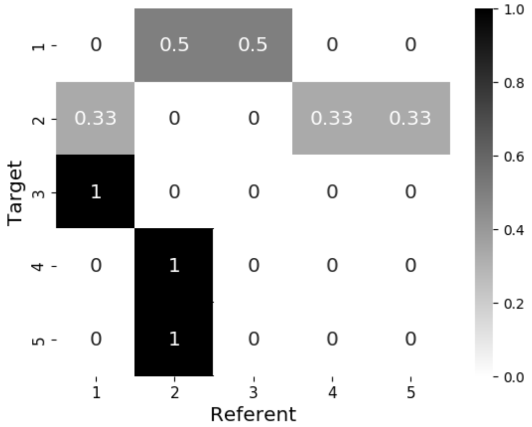


(b) Gold-Standard Weights

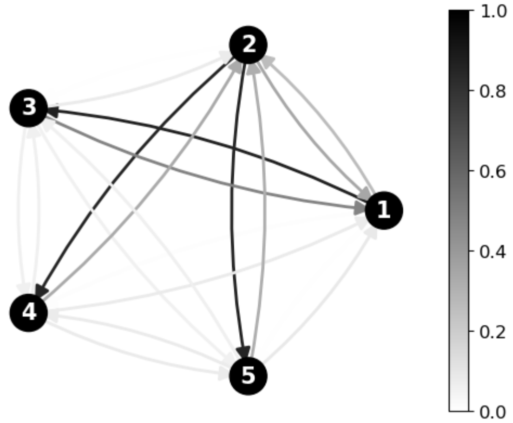


(c) Proximal Followers

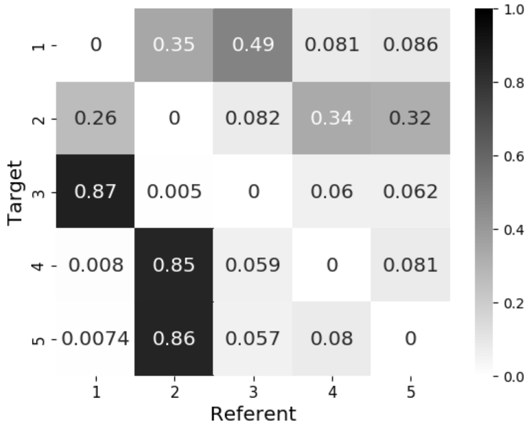


(d) Proximal Followers

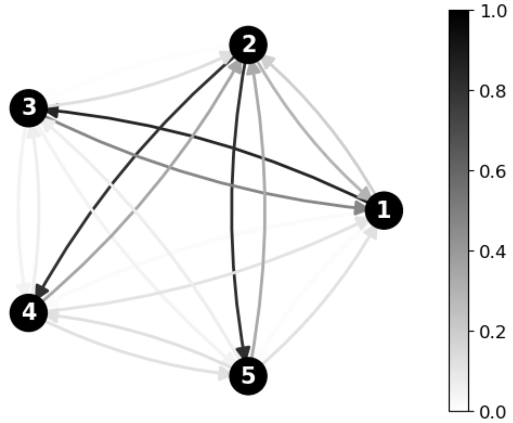


(e) Distant Followers

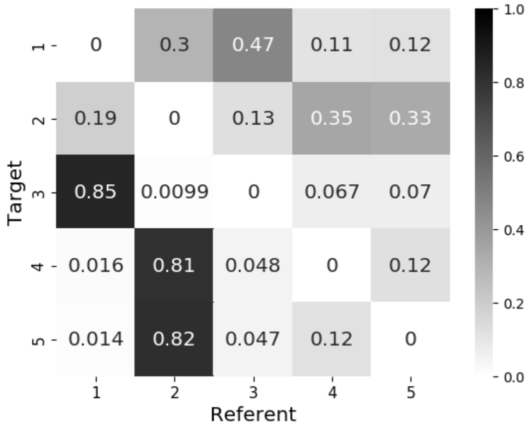


(f) Distant Followers

Figure S1.6: Gold-standard and average predicted attention weights for the random graph in Figure S1.1d. The KL divergence between gold-standard and predicted attention weights was 0.1477 for proximal followers ($p < 0.00001$) and 0.2054 for distant followers ($p = 0.00001$).

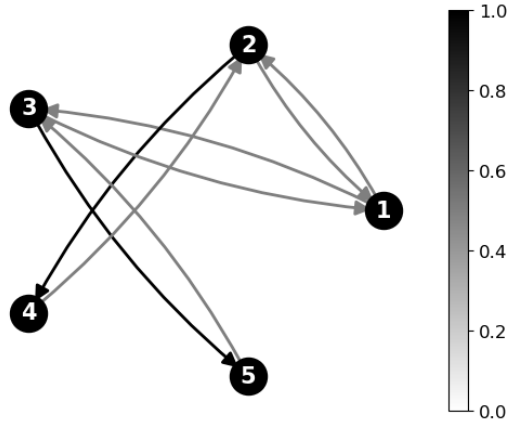


(a) Gold-Standard Weights

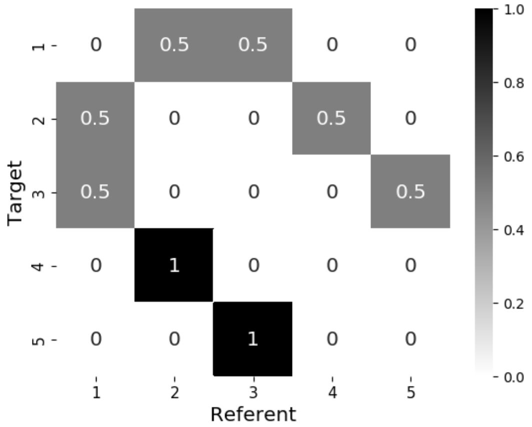


(b) Gold-Standard Weights

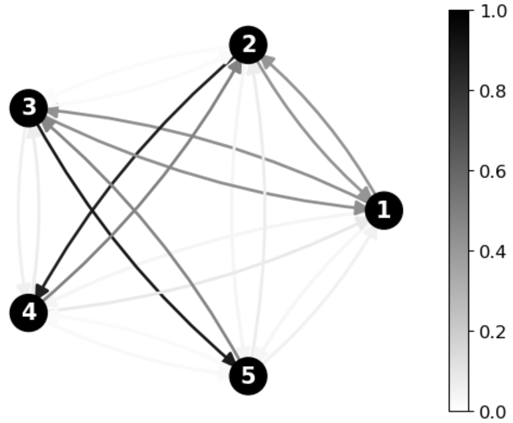


(c) Proximal Followers

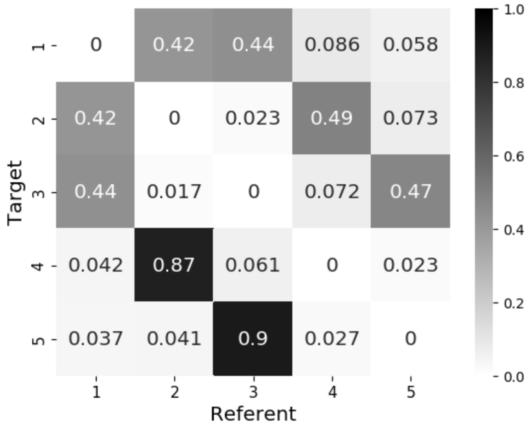


(d) Proximal Followers

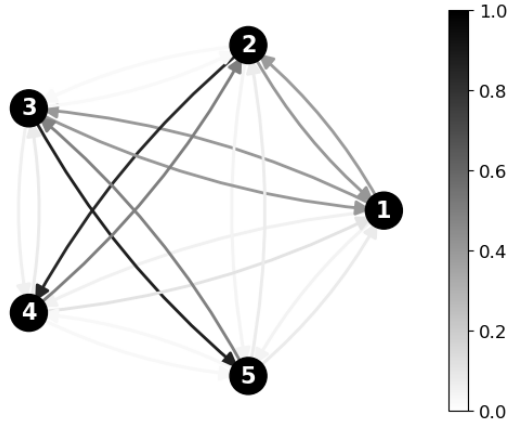


(e) Distant Followers

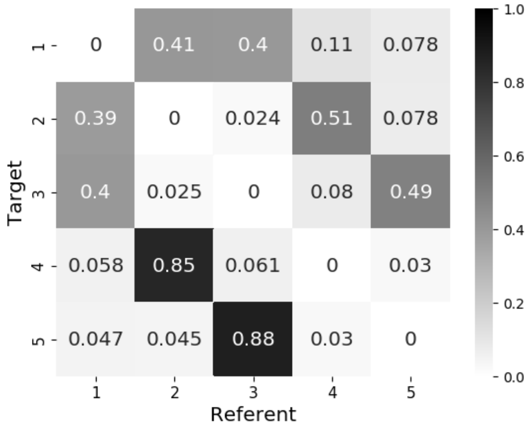


(f) Distant Followers

Figure S1.7: Gold-standard and average predicted attention weights for the random graph in Figure S1.1e. The KL divergence between gold-standard and predicted attention weights was 0.1198 for proximal followers ($p < 0.00001$) and 0.1465 for distant followers ($p < 0.00001$).

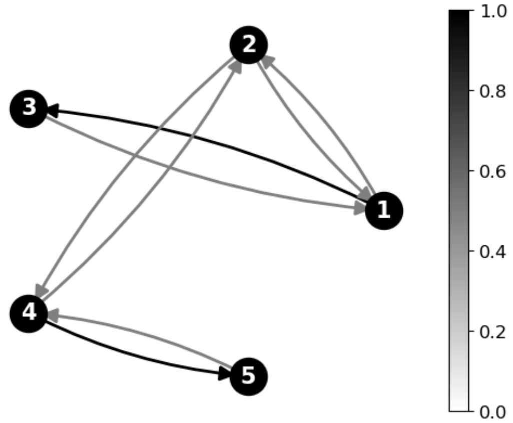


(a) Gold-Standard Weights

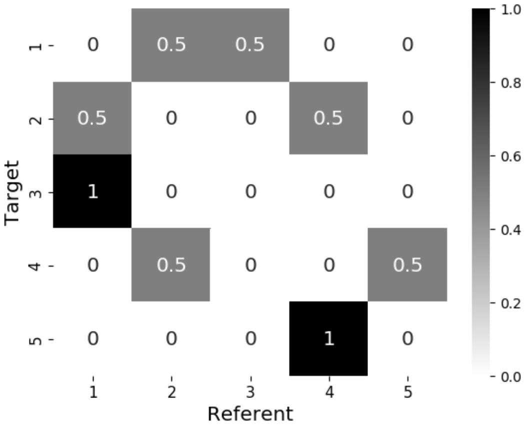


(b) Gold-Standard Weights

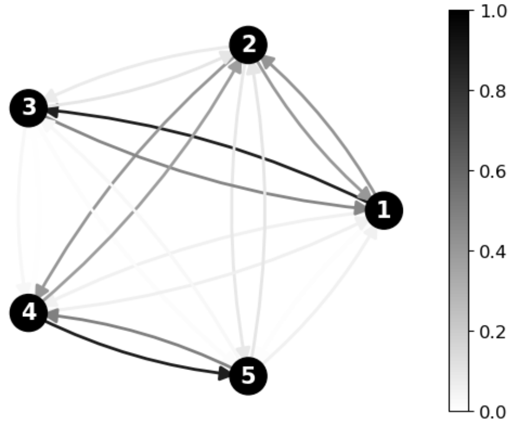


(c) Proximal Followers

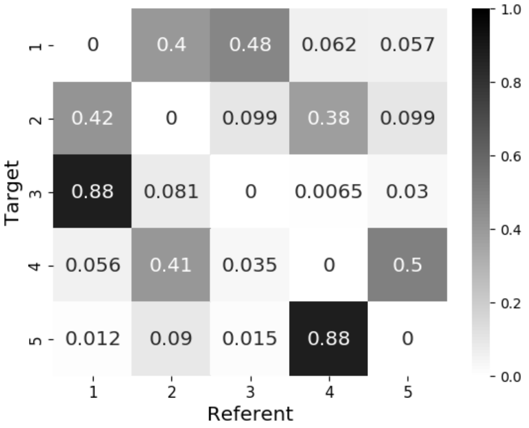


(d) Proximal Followers

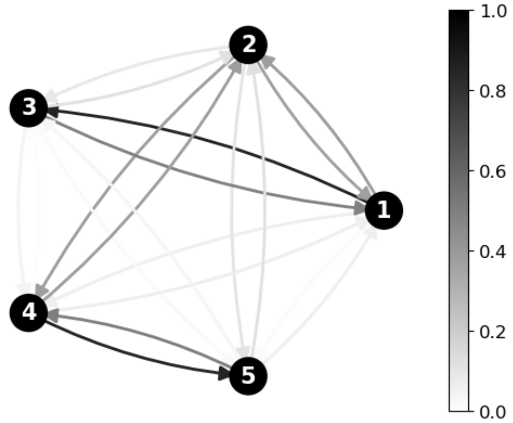


(e) Distant Followers

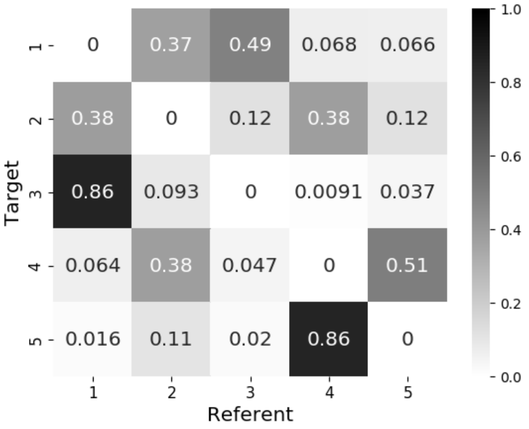


(f) Distant Followers

Figure S1.8: Gold-standard and average predicted attention weights for the random graph in Figure S1.1f. The KL divergence between gold-standard and predicted attention weights was 0.1407 for proximal followers ($p < 0.00001$) and 0.1710 for distant followers ($p < 0.00001$).

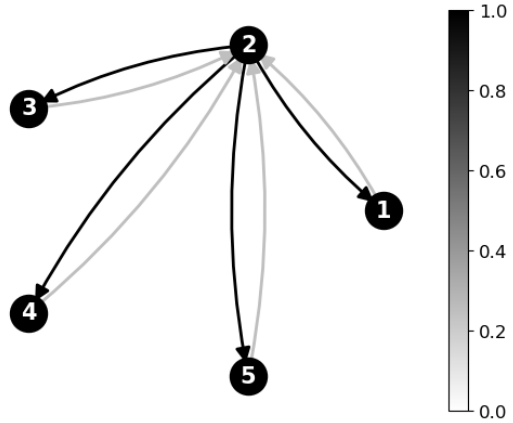


(a) Gold-Standard Weights

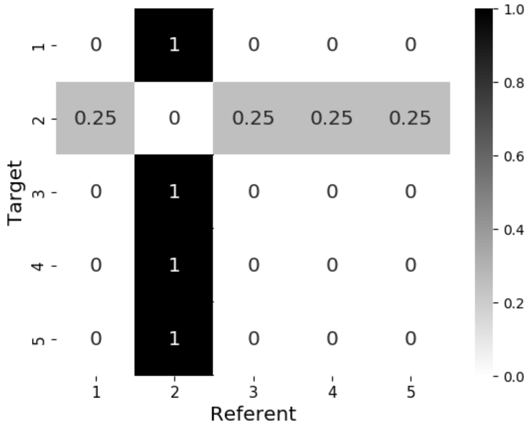


(b) Gold-Standard Weights

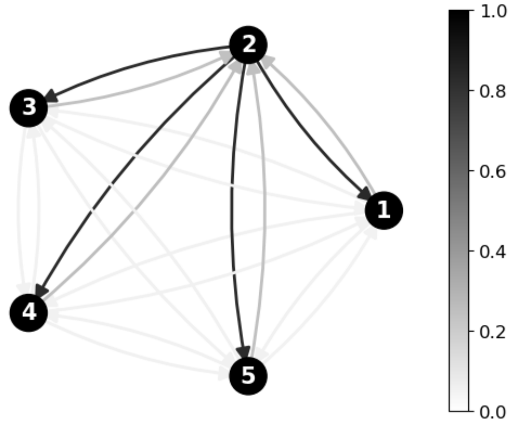


(c) Proximal Followers

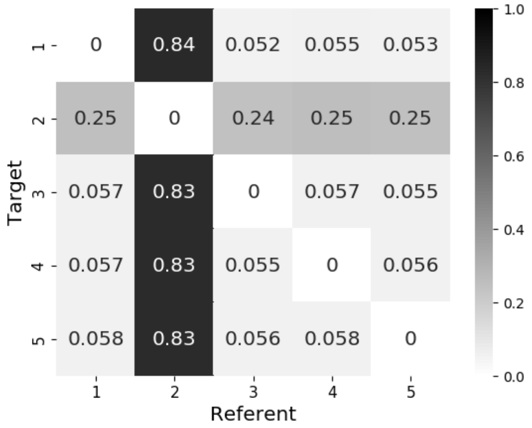


(d) Proximal Followers

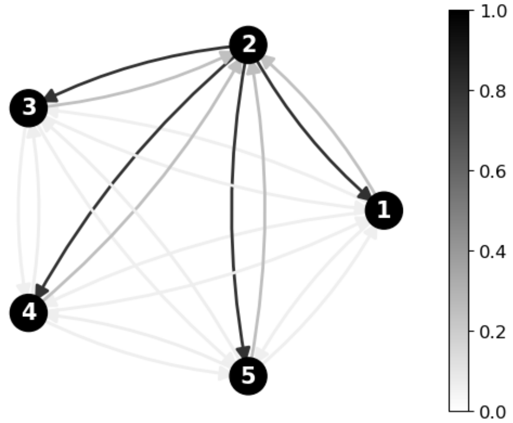


(e) Distant Followers

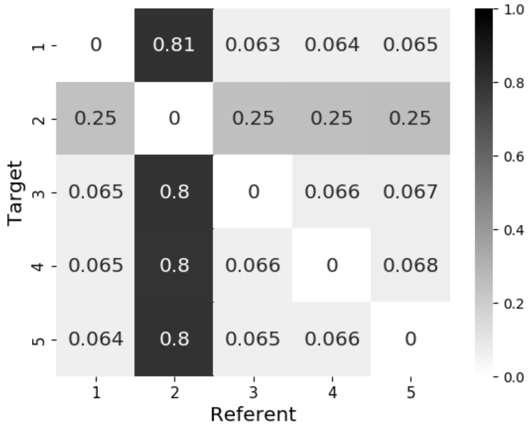


(f) Distant Followers

Figure S1.9: Gold-standard and average predicted attention weights for the random graph in Figure S1.1g. The KL divergence between gold-standard and predicted attention weights was 0.1467 for proximal followers ($p < 0.00001$) and 0.1750 for distant followers ($p < 0.00001$).

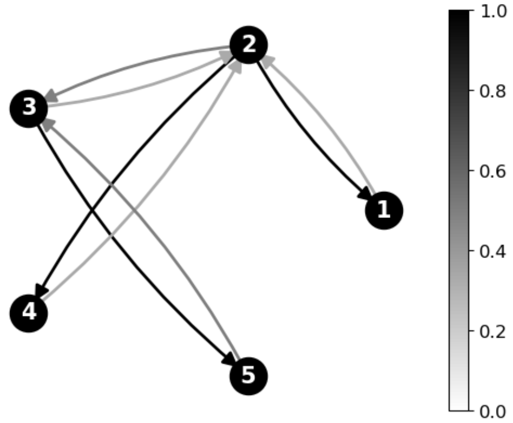


(a) Gold-Standard Weights

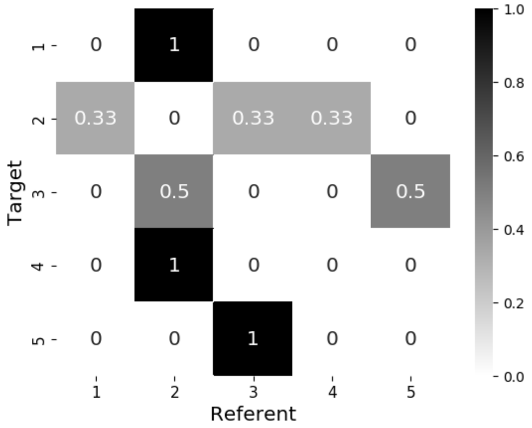


(b) Gold-Standard Weights

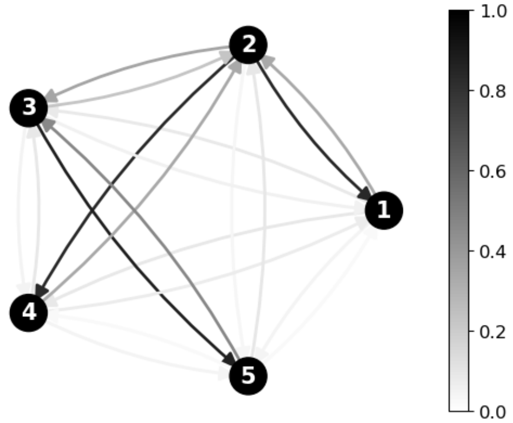


(c) Proximal Followers

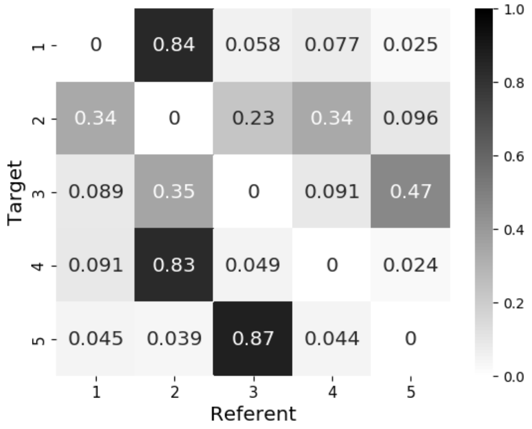


(d) Proximal Followers

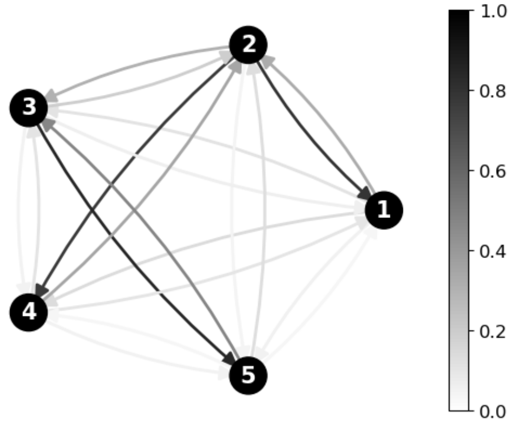


(e) Distant Followers

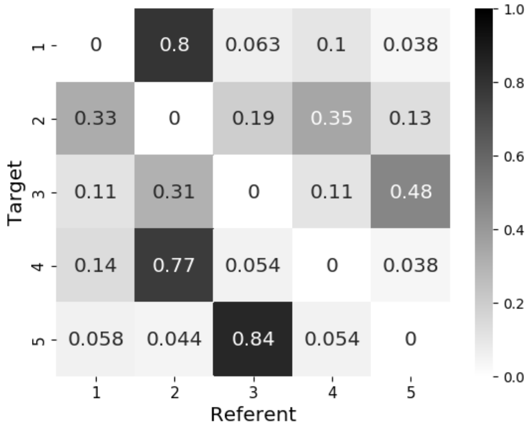


(f) Distant Followers

Figure S1.10: Gold-standard and average predicted attention weights for the random graph in Figure S1.1h. The KL divergence between gold-standard and predicted attention weights was 0.1634 for proximal followers ($p < 0.00001$) and 0.2208 for distant followers ($p = 0.00002$).

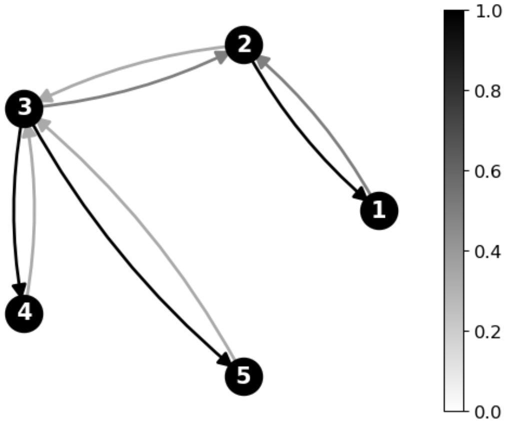


(a) Gold-Standard Weights

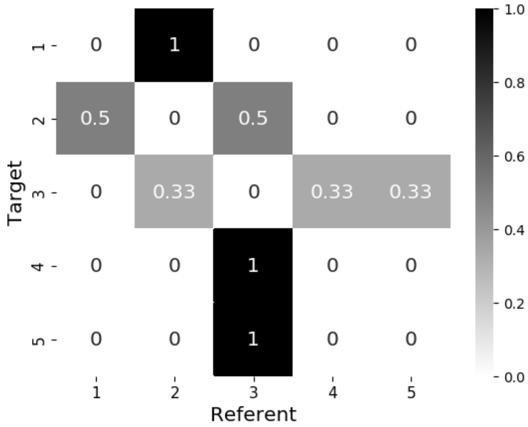


(b) Gold-Standard Weights

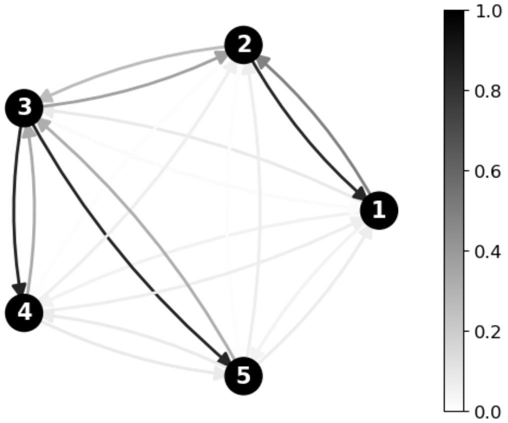


(c) Proximal Followers

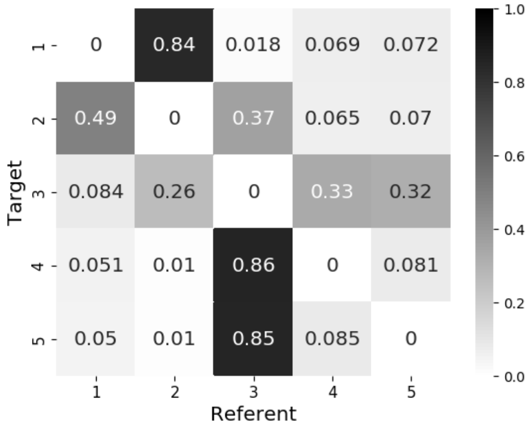


(d) Proximal Followers

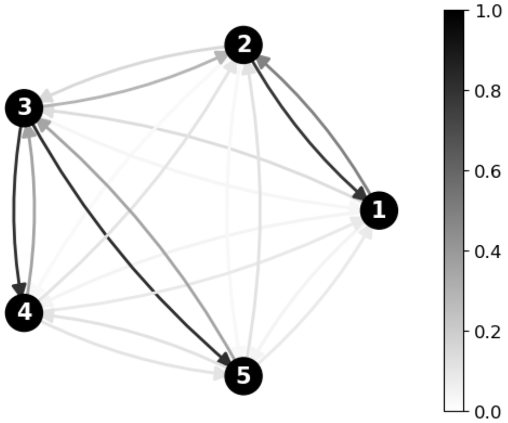


(e) Distant Followers

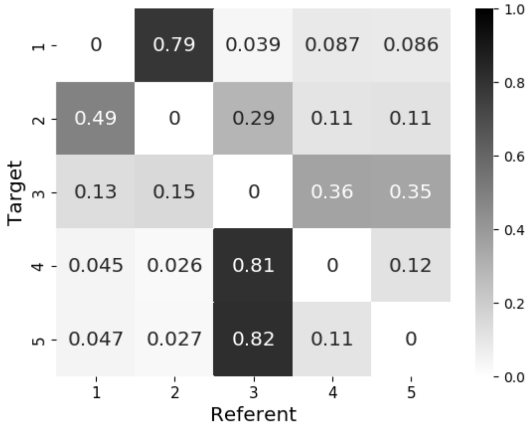


(f) Distant Followers

Figure S1.11: Gold-standard and average predicted attention weights for the random graph in Figure S1.1i. The KL divergence between gold-standard and predicted attention weights was 0.1465 for proximal followers ($p < 0.00001$) and 0.2297 for distant followers ($p < 0.00001$).

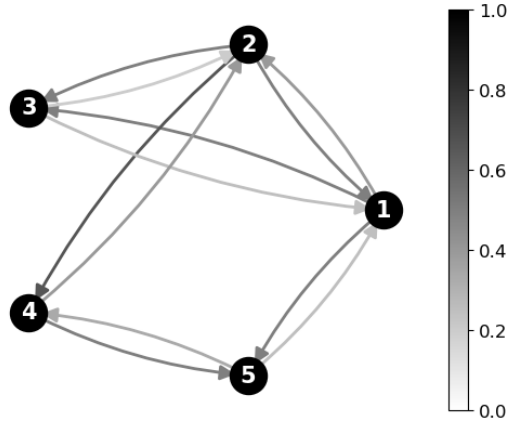


(a) Gold-Standard Weights

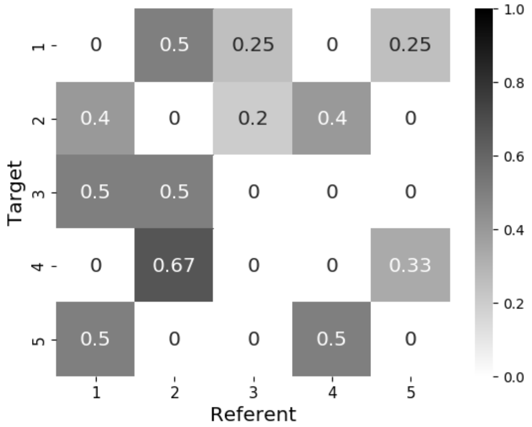


(b) Gold-Standard Weights

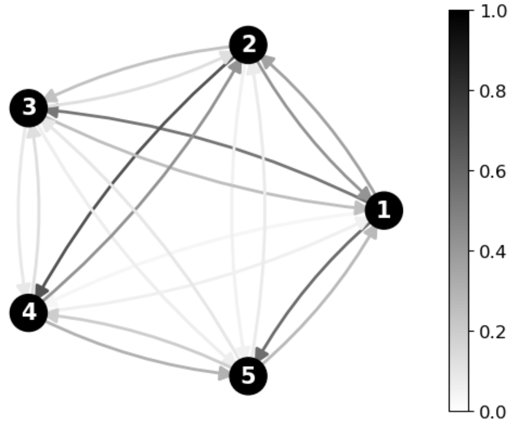


(c) Proximal Followers

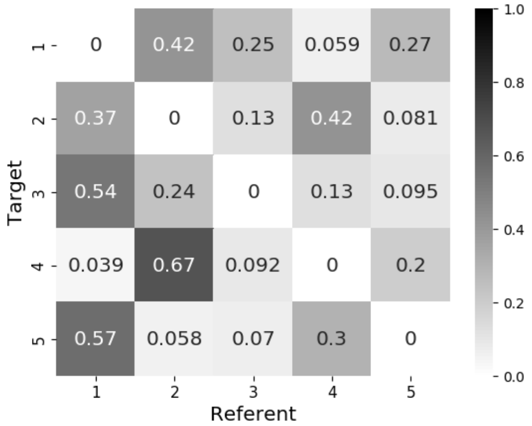


(d) Proximal Followers

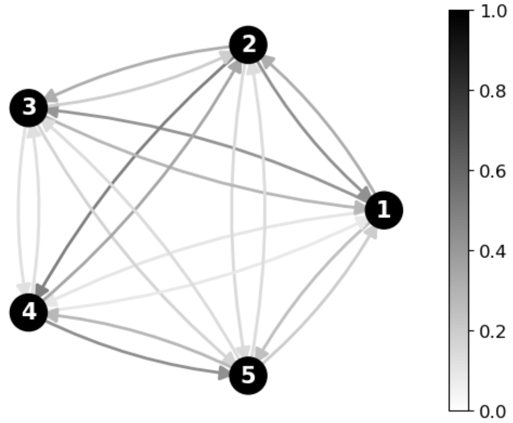


(e) Distant Followers

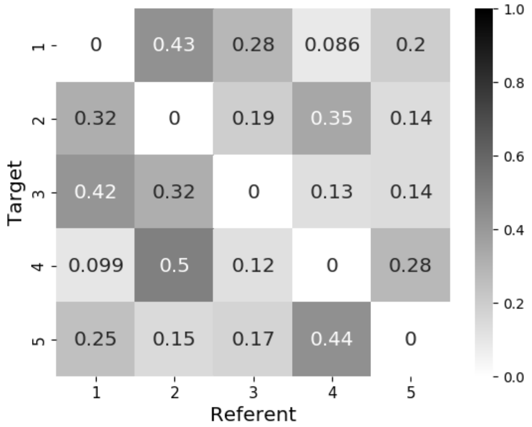


(f) Distant Followers

Figure S1.12: Gold-standard and average predicted attention weights for the random graph in Figure S1.2a. The KL divergence between gold-standard and predicted attention weights was 0.1711 for proximal followers ($p < 0.00001$) and 0.2474 for distant followers ($p = 0.00013$).

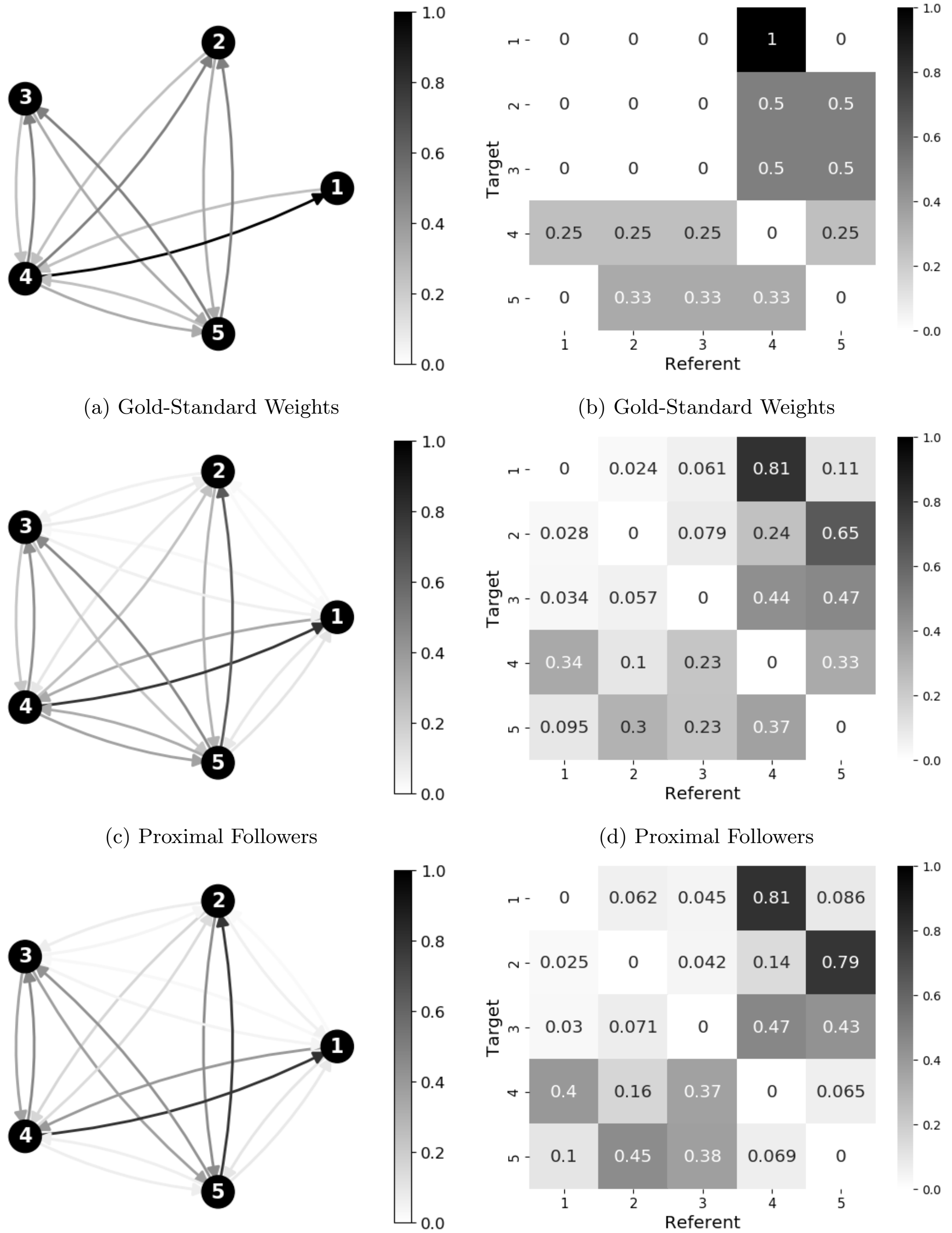


(a) Gold-Standard Weights (b) Gold-Standard Weights

(c) Proximal Followers (d) Proximal Followers

(e) Distant Followers (f) Distant Followers

Figure S1.13: Gold-standard and average predicted attention weights for the random graph in Figure S1.2b. The KL divergence between gold-standard and predicted attention weights was 0.1524 for proximal followers ($p = 0.00001$) and 0.2674 for distant followers ($p = 0.00053$).

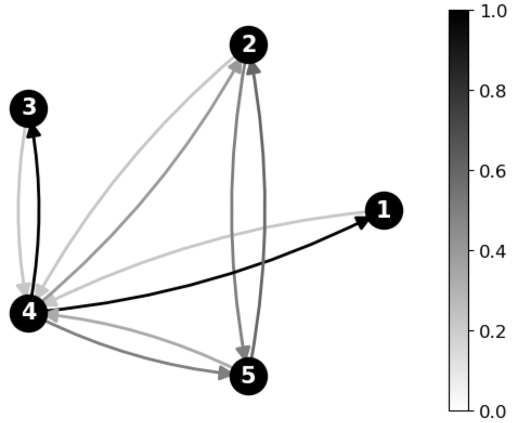


(a) Gold-Standard Weights

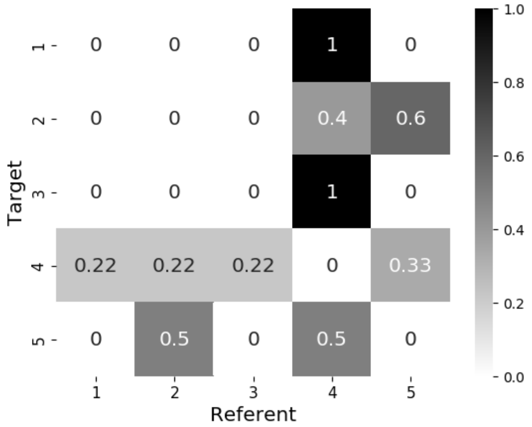


(b) Gold-Standard Weights

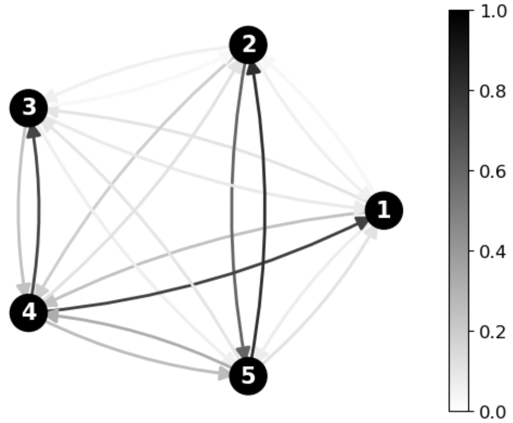


(c) Proximal Followers

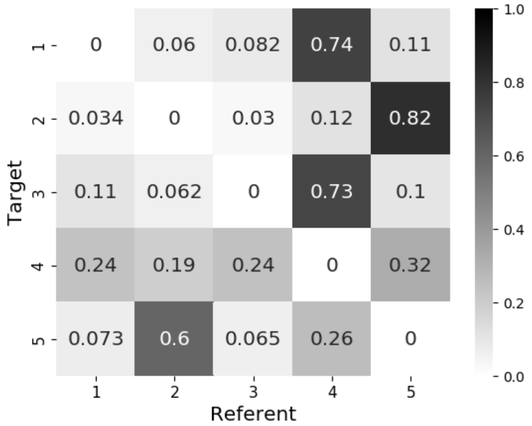


(d) Proximal Followers

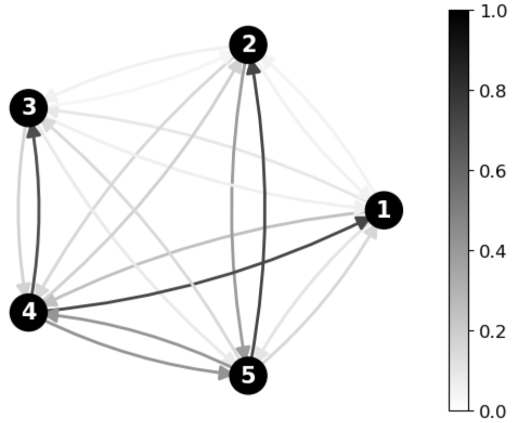


(e) Distant Followers

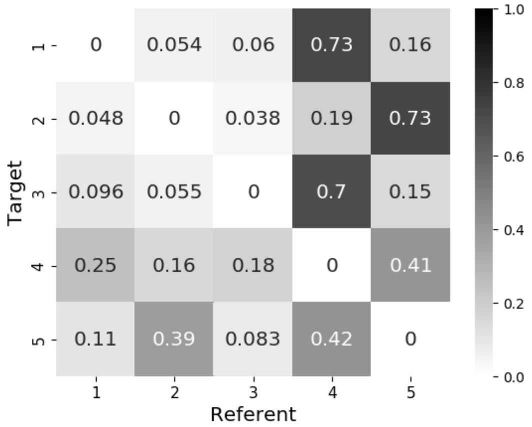


(f) Distant Followers

Figure S1.14: Gold-standard and average predicted attention weights for the random graph in Figure S1.2c. The KL divergence between gold-standard and predicted attention weights was 0.2322 for proximal followers ($p = 0.00001$) and 0.2211 for distant followers ($p = 0.00001$).

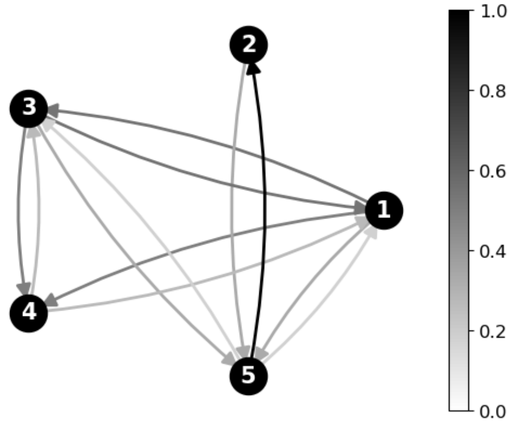


(a) Gold-Standard Weights

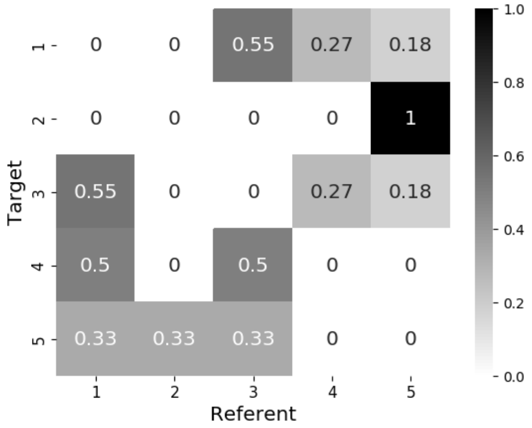


(b) Gold-Standard Weights

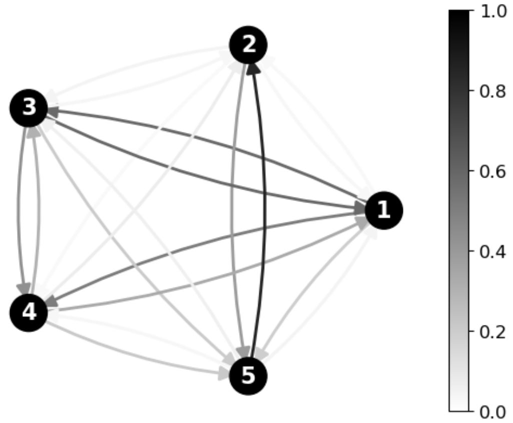


(c) Proximal Followers

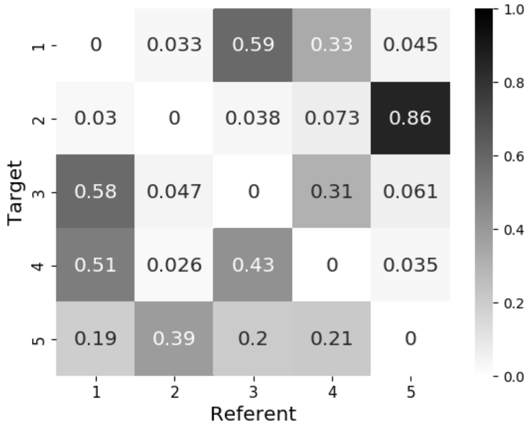


(d) Proximal Followers

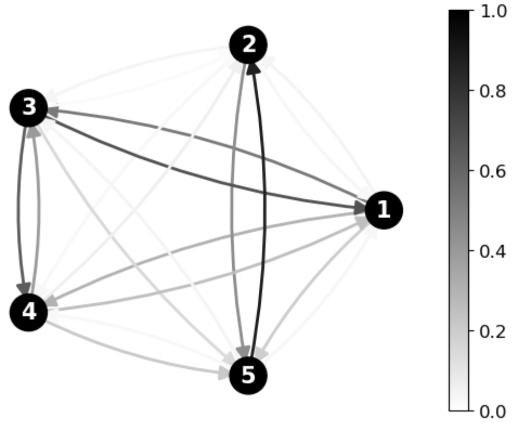


(e) Distant Followers

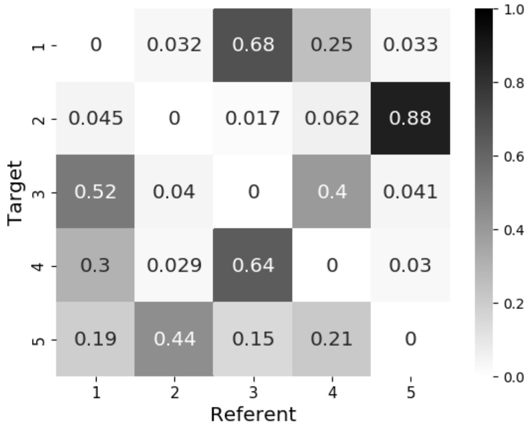


(f) Distant Followers

Figure S1.15: Gold-standard and average predicted attention weights for the random graph in Figure S1.2d. The KL divergence between gold-standard and predicted attention weights was 0.1594 for proximal followers ($p = 0.00003$) and 0.2024 for distant followers ($p = 0.00018$).

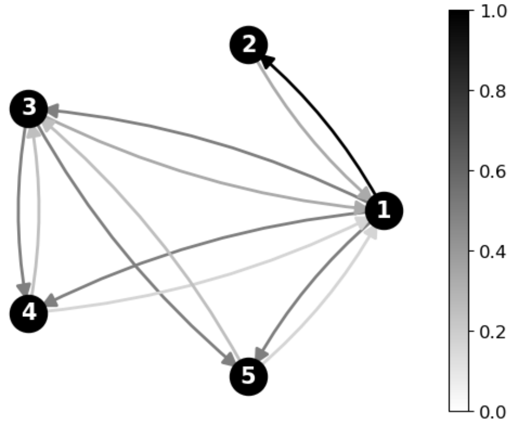


(a) Gold-Standard Weights

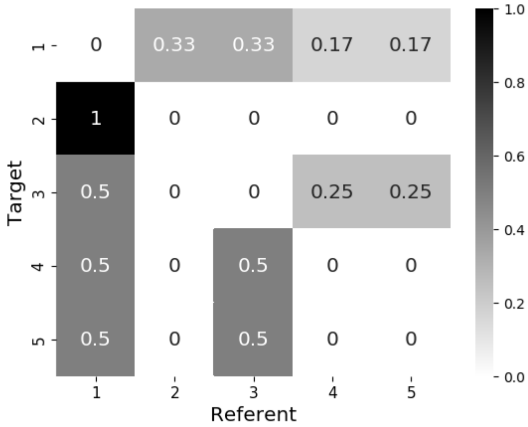


(b) Gold-Standard Weights

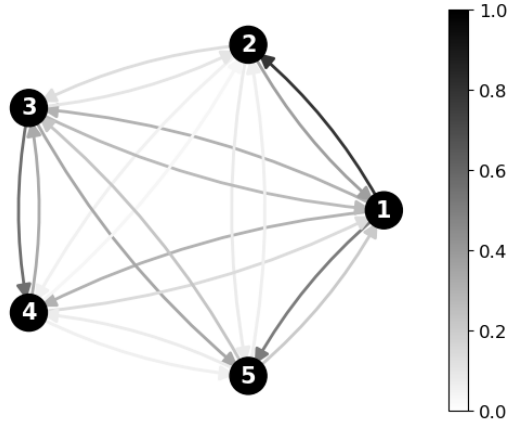

(c) Proximal Followers

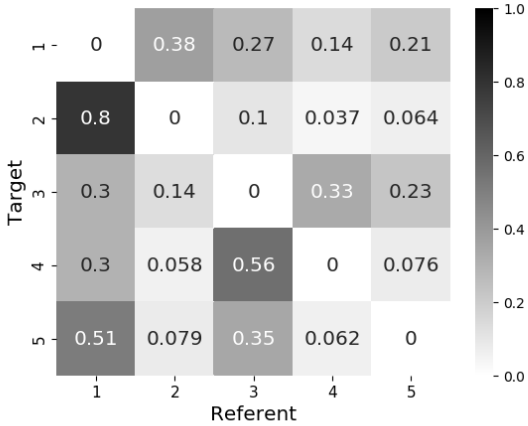


(d) Proximal Followers

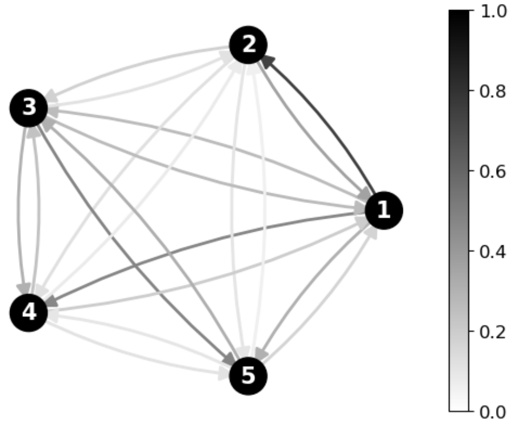

(e) Distant Followers

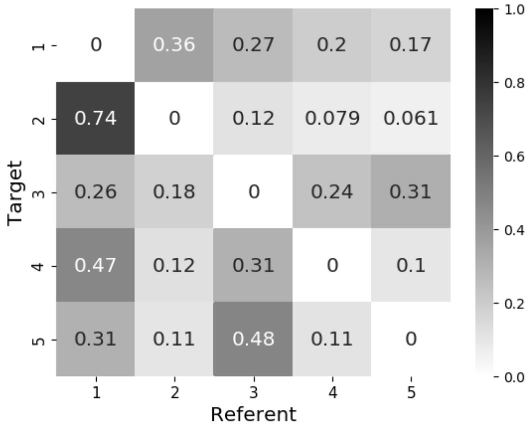


(f) Distant Followers

Figure S1.16: Gold-standard and average predicted attention weights for the random graph in Figure S1.2e. The KL divergence between gold-standard and predicted attention weights was 0.1620 for proximal followers ($p = 0.00002$) and 0.2249 for distant followers ($p = 0.00026$).

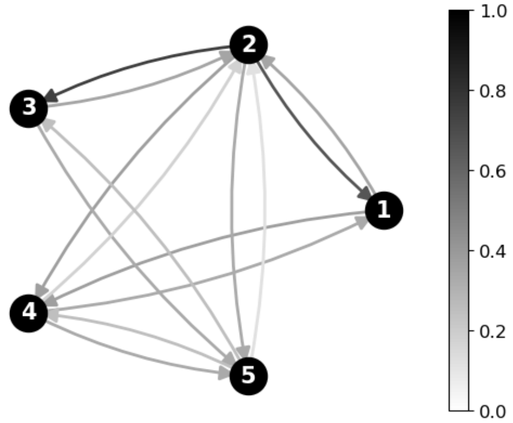


(a) Gold-Standard Weights

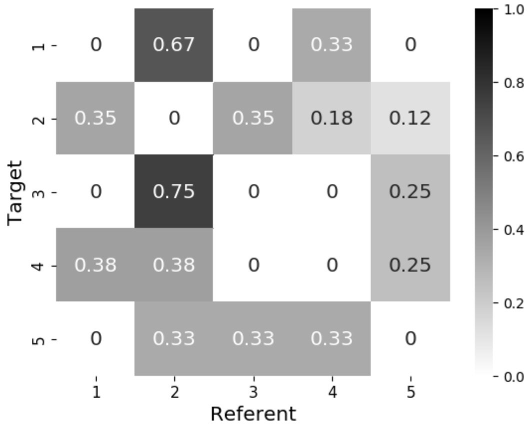


(b) Gold-Standard Weights

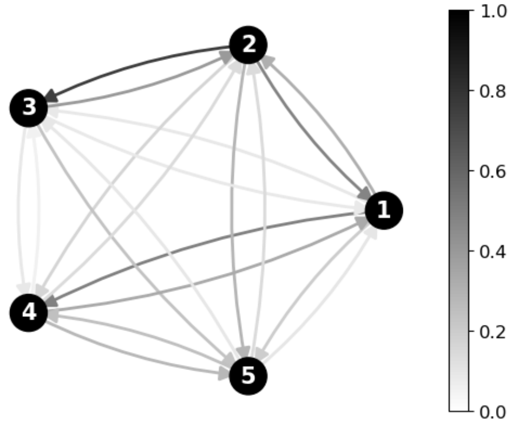


(c) Proximal Followers

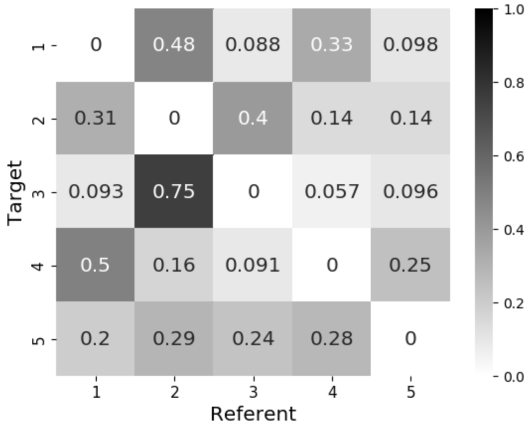


(d) Proximal Followers

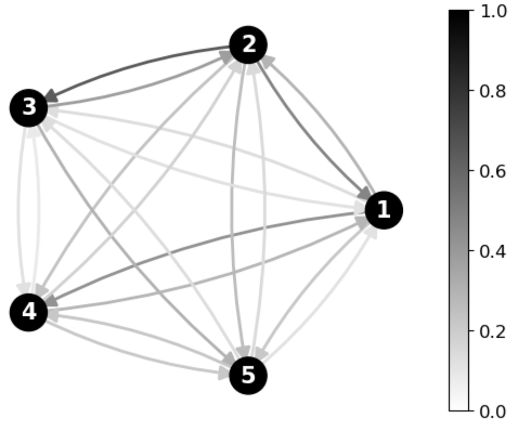


(e) Distant Followers

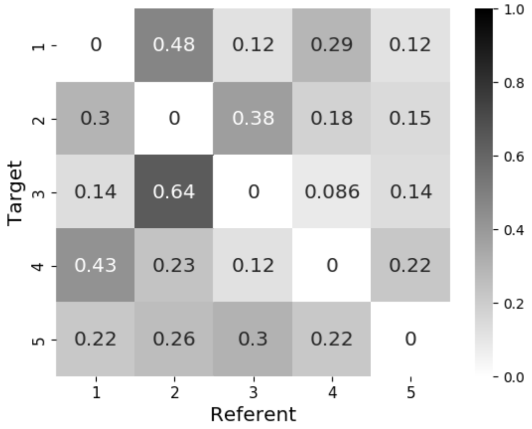


(f) Distant Followers

Figure S1.17: Gold-standard and average predicted attention weights for the random graph in Figure S1.2f. The KL divergence between gold-standard and predicted attention weights was 0.1785 for proximal followers ($p = 0.00148$) and 0.0779 for distant followers ($p = 0.00195$).

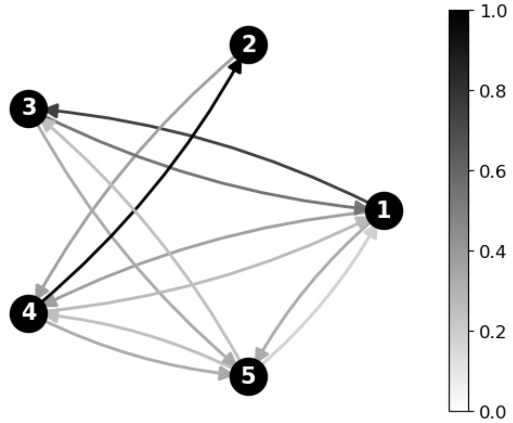


(a) Gold-Standard Weights

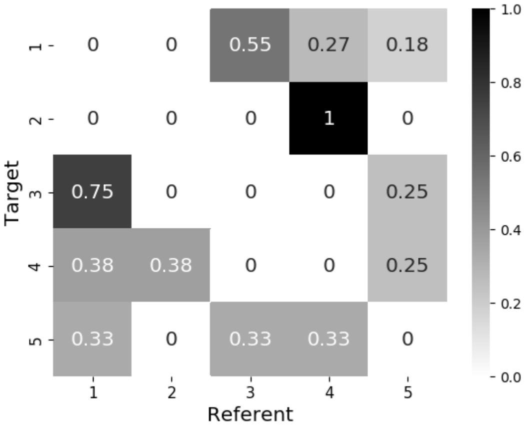


(b) Gold-Standard Weights

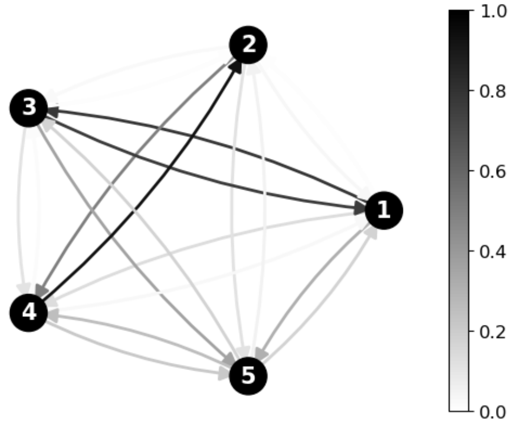


(c) Proximal Followers

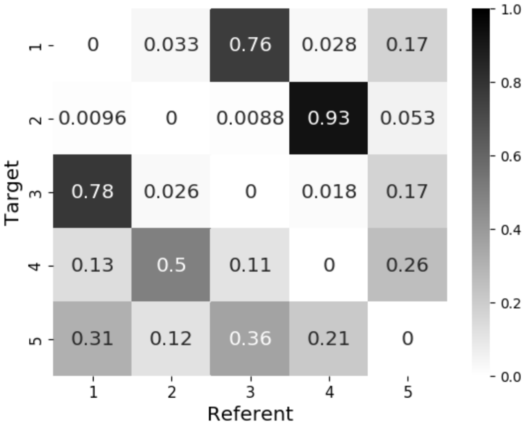


(d) Proximal Followers

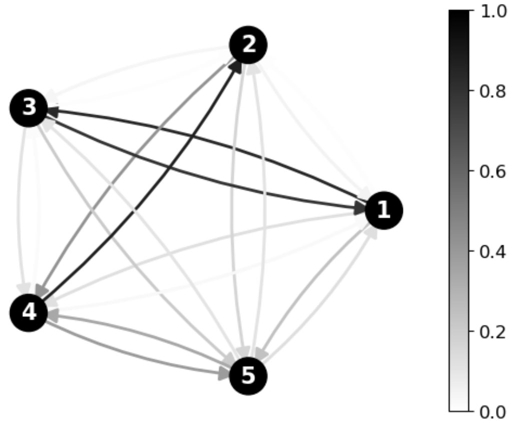


(e) Distant Followers

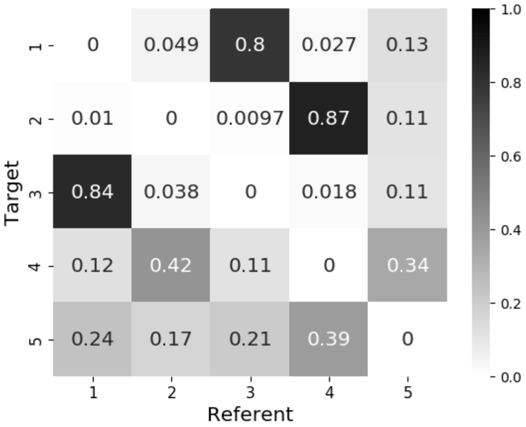


(f) Distant Followers

Figure S1.18: Gold-standard and average predicted attention weights for the random graph in Figure S1.2g. The KL divergence between gold-standard and predicted attention weights was 0.2009 for proximal followers ($p < 0.00001$) and 0.2539 for distant followers ($p = 0.00010$).

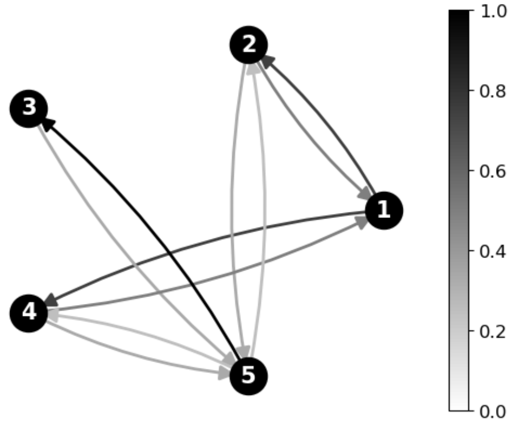


(a) Gold-Standard Weights

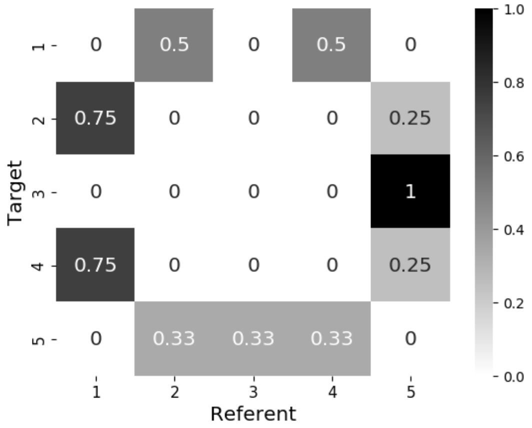


(b) Gold-Standard Weights

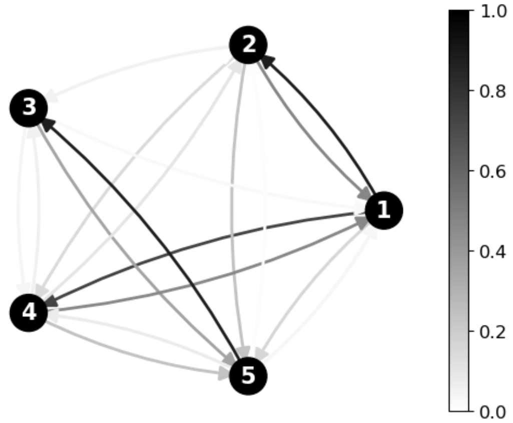

(c) Proximal Followers

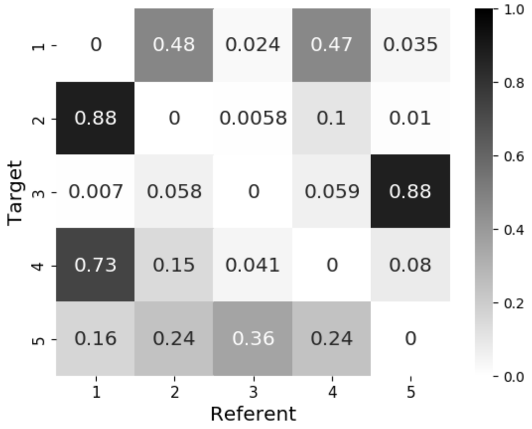


(d) Proximal Followers

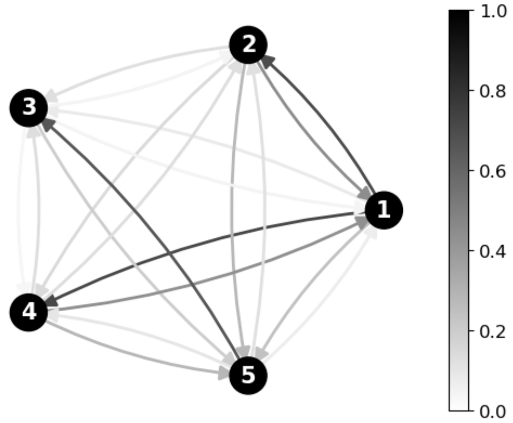

(e) Distant Followers

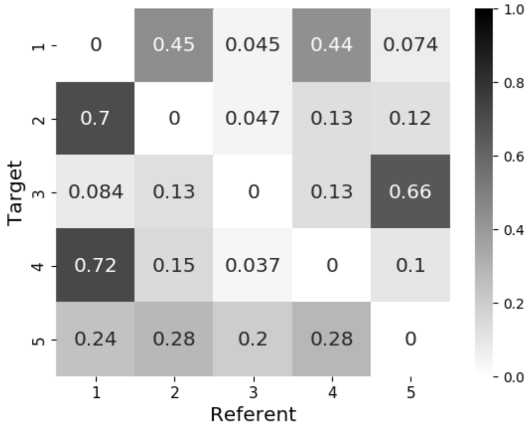


(f) Distant Followers

Figure S1.19: Gold-standard and average predicted attention weights for the random graph in Figure S1.2h. The KL divergence between gold-standard and predicted attention weights was 0.2750 for proximal followers ($p = 0.00023$) and 0.2646 for distant followers ($p = 0.00017$).

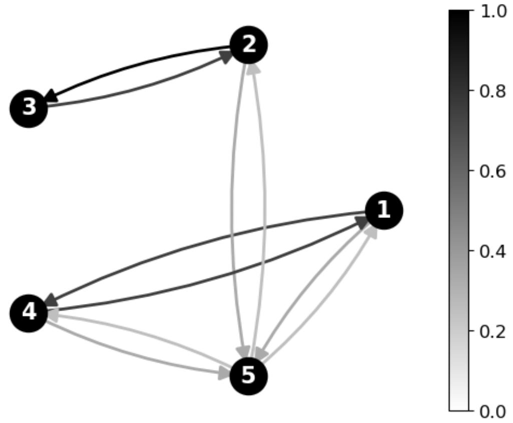


(a) Gold-Standard Weights

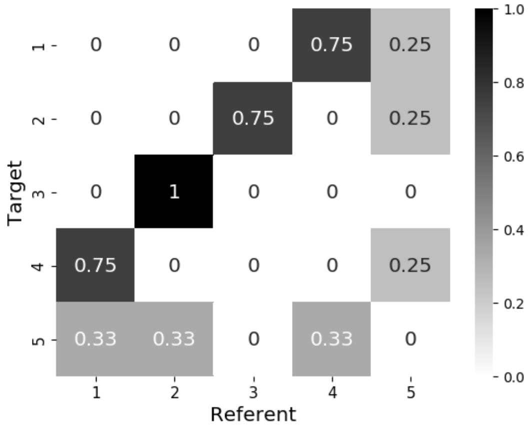


(b) Gold-Standard Weights

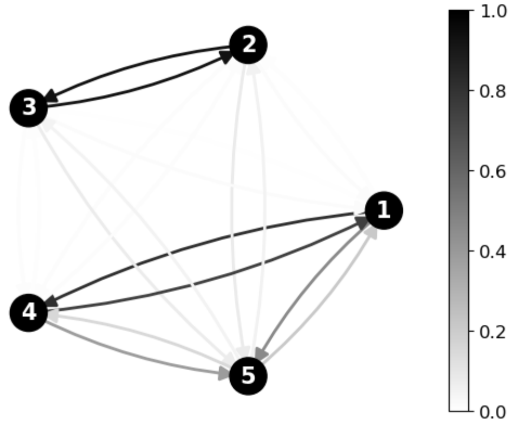


(c) Proximal Followers

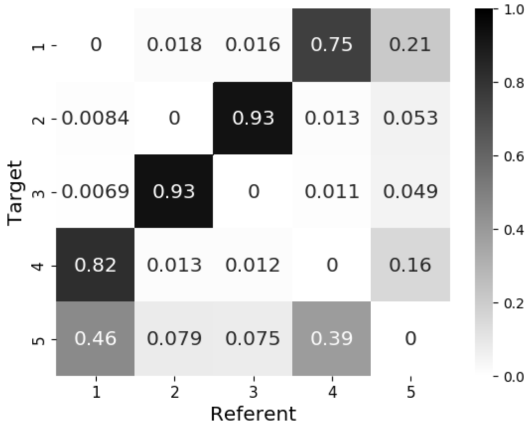


(d) Proximal Followers

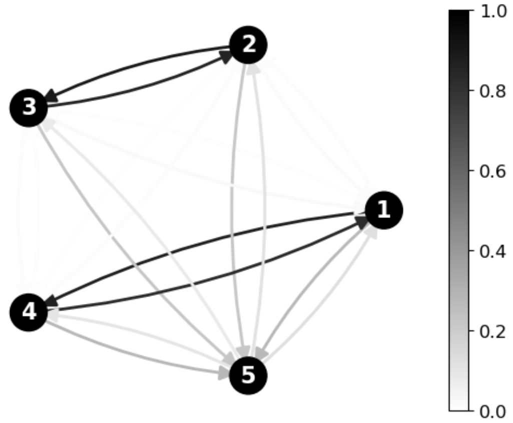


(e) Distant Followers

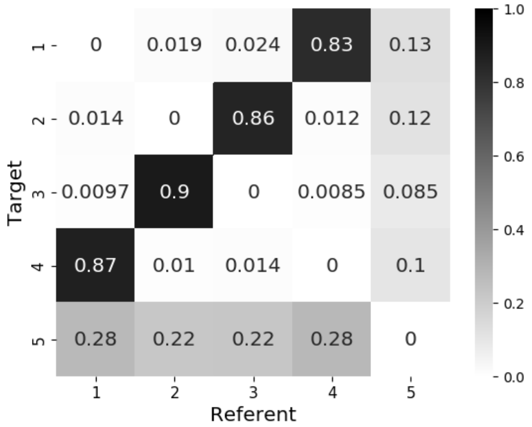


(f) Distant Followers

Figure S1.20: Gold-standard and average predicted attention weights for the random graph in Figure S1.2i. The KL divergence between gold-standard and predicted attention weights was 0.1428 for proximal followers ($p < 0.00001$) and 0.1319 for distant followers ($p < 0.00001$).

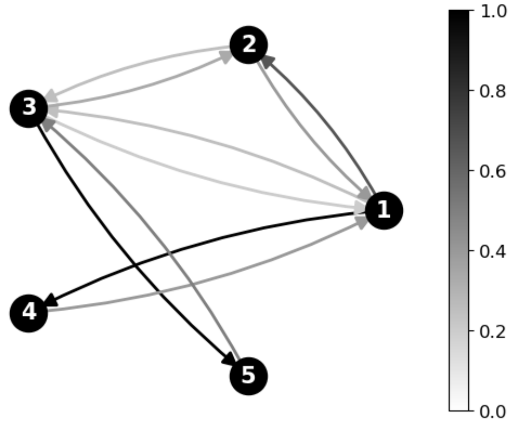


(a) Gold-Standard Weights

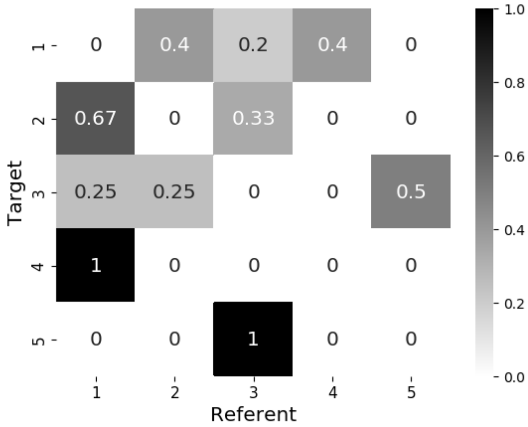


(b) Gold-Standard Weights

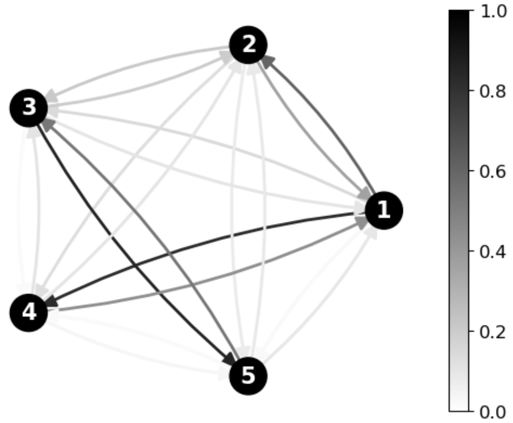


(c) Proximal Followers

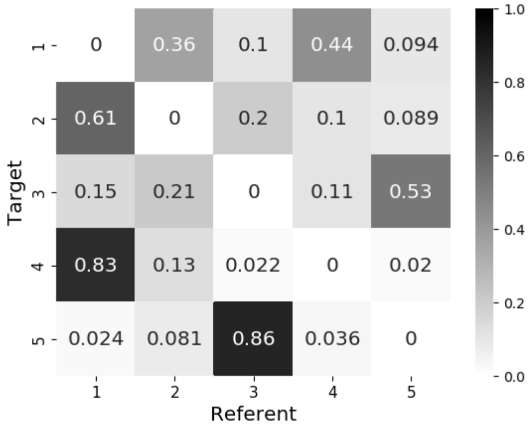


(d) Proximal Followers

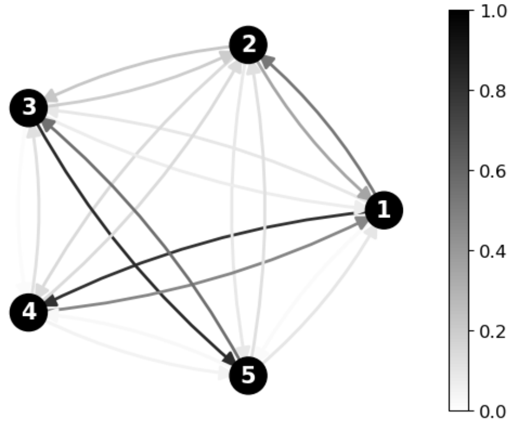


(e) Distant Followers

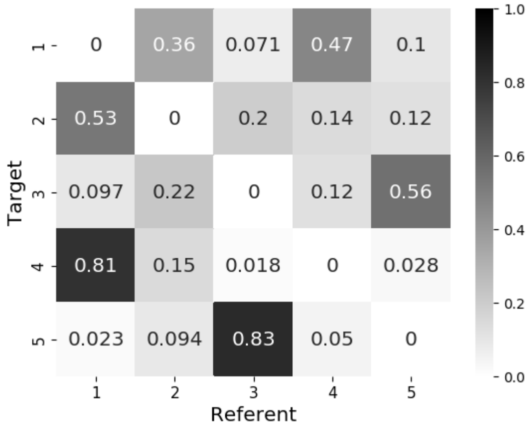


(f) Distant Followers

Figure S1.21: Gold-standard and average predicted attention weights for the random graph in Figure S1.2j. The KL divergence between gold-standard and predicted attention weights was 0.1694 for proximal followers ($p < 0.00001$) and 0.2227 for distant followers ($p = 0.00001$).

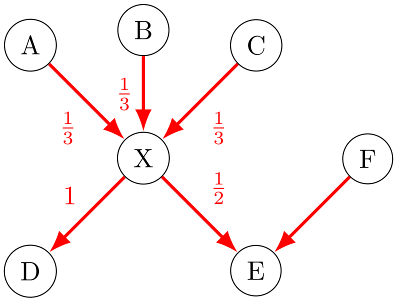


Figure S2.1: Toy example illustrating the computation of the gold-standard weights. The rational numbers represent the probability that the corresponding arrows will be used in the generative process. The gold-standard weights are their normalized values.

46 X and E is only used once every two times. The gold-standard attention weights normalize these generative
47 probability; for example,

$$(w_{\mathrm{X,A}}, w_{\mathrm{X,B}}, w_{\mathrm{X,C}}, w_{\mathrm{X,D}}, w_{\mathrm{X,E}}) = \frac{(\frac{1}{3}, \frac{1}{3}, \frac{1}{3}, 1, \frac{1}{2})}{\frac{1}{3} + \frac{1}{3} + \frac{1}{3} + 1 + \frac{1}{2}} \approx (0.133, 0.133, 0.133, 0.400, 0.200) \quad \text{(S2.1)}$$

48 in Figure S2.1.
